# Sensory integration of food availability and population density during the diapause exit decision involves insulin-like signaling in *Caenorhabditis elegans*

**DOI:** 10.1101/2024.03.20.586022

**Authors:** Mark G Zhang, Maedeh Seyedolmohadesin, Soraya Hawk, Heenam Park, Nerissa Finnen, Frank Schroeder, Vivek Venkatachalam, Paul W Sternberg

**Affiliations:** Division of Biology and Biological Engineering, California Institute of Technology, Pasadena, CA, USA; Physics Department, Northeastern University, Boston, MA, USA; Boyce Thompson Institute and Department of Chemistry and Chemical Biology, Cornell University, Ithaca, NY, USA

**Keywords:** dauer, diapause, decision, sensory, integration, insulin, neuropeptide

## Abstract

Decisions made over long time scales, such as life cycle decisions, require coordinated interplay between sensory perception and sustained gene expression. The *Caenorhabditis elegans* dauer (or diapause) exit developmental decision requires sensory integration of population density and food availability to induce an all-or-nothing organismal-wide response, but the mechanism by which this occurs remains unknown. Here, we demonstrate how the ASJ chemosensory neurons, known to be critical for dauer exit, perform sensory integration at both the levels of gene expression and calcium activity. In response to favorable conditions, dauers rapidly produce and secrete the dauer exit-promoting insulin-like peptide INS-6. Expression of *ins-6* in the ASJ neurons integrate population density and food level and can reflect decision commitment since dauers committed to exiting have higher *ins-6* expression levels than those of non-committed dauers. Calcium imaging in dauers reveals that the ASJ neurons are activated by food, and this activity is suppressed by pheromone, indicating that sensory integration also occurs at the level of calcium transients. We find that *ins-6* expression in the ASJ neurons depends on neuronal activity in the ASJs, cGMP signaling, a CaM-kinase pathway, and the pheromone components ascr#8 and ascr#2. We propose a model in which decision commitment to exit the dauer state involves an autoregulatory feedback loop in the ASJ neurons that promotes high INS-6 production and secretion. These results collectively demonstrate how insulin-like peptide signaling helps animals compute long-term decisions by bridging sensory perception to decision execution.

**Summary/Significance Statement:** Animals must respond appropriately to multiple sensory stimuli to make informed decisions. It remains unclear how the nervous system is able to integrate different sensory cues and propagate that information towards making decisions over longer timescales. We use the nematode *Caenorhabditis elegans* to investigate how sensory integration occurs during the decision to exit diapause, a stress-resistant developmentally arrested state triggered by multiple sensory inputs including food availability and population density. We show how expression of an insulin-like peptide critical to dauer exit reflects the sensory integration process and decision commitment, and we dissect the regulation of this insulin-like peptide’s expression. Our study explicitly analyzes the relationship between neuronal activity and neuropeptide expression during a complex decision with diverse sensory inputs.

## Introduction

A fundamental goal of neuroscience is to understand how nervous systems sense, integrate, and interpret diverse stimuli to deliver the appropriate output. The duration over which these processes occur spans from shorter timescales, such as during reflexive responses to harsh stimuli or chemotaxis towards attractive stimuli (Browne et al., 2017; Gao, 2014; Hart, 2006; Paintal, 1961; Vriens et al., 2014), to longer timescales, such as migration or hibernation decisions in response to changing weather patterns, temperature, and food availability (Evans et al., 2016; Komal et al., 2017; Mohr et al., 2020; Ogonowski and Conway, 2009). We lack a clear understanding of how neuronal activity, which occurs over short timescales of milliseconds to seconds, informs downstream processes that occur over timescales of hours or longer. One such decision that takes place over a longer timescale is the developmental decision to enter (or exit) diapause, a temporarily suspended developmental state that protects against environmental stress and promotes dispersal. Diapause is evolutionarily conserved across metazoans and requires integration of environmental cues to inform a coordinated, organismal-wide decision (Denlinger et al., 2012; Hand et al., 2016; Podrabsky and Hand, 2015)

To better understand how a compact nervous system can interpret sensory signals to coordinate a organismal-wide decision that takes place over multiple hours, we studied the dauer exit decision. During early larval growth, *C. elegans* choose between two developmental fates depending on environmental conditions (**Fig. 1A**). Under favorable conditions, larvae undergo reproductive growth, whereas under unfavorable conditions, larvae enter the stress-resistant, long-lived developmentally arrested diapause state known as dauer (Cassada and Russell, 1975; Golden and Riddle, 1984, 1982). While in the dauer state, *C. elegans* continually assess their surroundings to detect environmental improvement; when conditions sufficiently improve, animals exit the dauer state and return to the reproductive cycle as late-stage larvae. Of the various environmental inputs to the dauer entry and exit decisions, the strongest input is a ratio of food to pheromone (Golden and Riddle, 1982). Here, “pheromone” refers to a mixture of secreted dauer-regulating signaling molecules, termed ascarosides, that collectively convey population density (Ludewig and Schroeder, 2018; McGrath and Ruvinsky, 2019).

**Figure 1.**
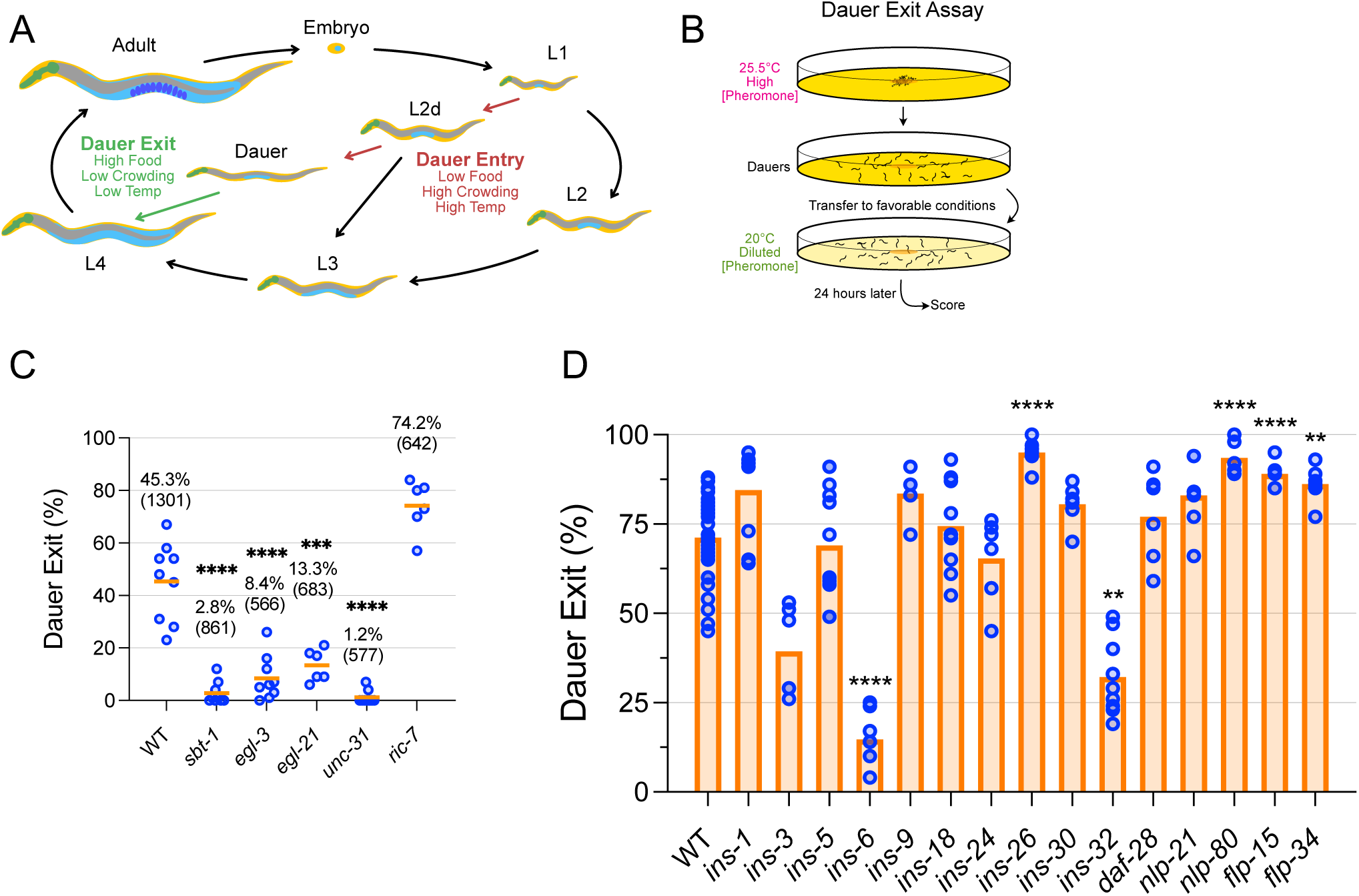
Neuropeptides, especially the insulin-like peptide *ins-6*, are critical for dauer exit. (A) During development, *C. elegans* makes multiple developmental decisions including whether to enter and when to exit the developmentally arrested state called dauer. This decision depends mainly on food availability, crowding, and temperature. (B) Overview of the dauer exit assay. Animals were induced to become dauer via growth on conditions of high pheromone concentration and high temperature. Non-dauers were removed using SDS and surviving dauers were transferred to conditions of intermediate pheromone concentrations and lower temperature to stimulate between 40-80% of dauers to exit after 24 hours. (C) Dauer exit rates of mutants defective for neuropeptide processing (*sbt-1*, *egl-3*, *egl-21*) or signaling (*unc-31*, *ric-7*) compared to that of wild type (WT). Means are written and shown by the orange line, and total number of animals scored is indicated in parentheses. (D) Dauer exit rates of mutants defective for single neuropeptide genes. Data compiled from multiple independent experiments, and statistical analyses were only performed between mutant and wild-type samples measured in the same experiment. Bars indicate means. See Figure S1 for analysis of individual mutants. For (C) and (D), each dot is the dauer exit % from an assay plate containing 50-100 animals each. ****, p<0.0001, ***, p<0.001, **, p<0.01, compared to “WT” by Welch ANOVA with Dunnett’s T3 multiple comparison correction. Comparisons to “WT” were not statistically significant unless indicated otherwise.

The genetic pathways and neuroendocrine signaling mechanisms that govern the dauer entry and exit decisions (reviewed in Androwski et al., 2017; Fielenbach and Antebi, 2008; Hu, 2007) including a cGMP pathway, a TGF-β-like pathway, an insulin/insulin-like growth factor (IGF)-1 signaling (IIS) pathway, and a steroid hormone pathway. While considerably more attention has been paid towards the dauer entry decision, previous studies identified two insulin-like peptides, INS-6 and DAF-28, as regulators of dauer exit that work within the IIS pathway (Cornils et al., 2011; Fernandes de Abreu et al., 2014).

Despite the wealth of knowledge collected on the *C. elegans* dauer entry and exit decisions, many fundamental questions remain, including: (1) How does sensory integration of food, pheromone, and temperature (amongst other possible cues) occur? and (2) How does short-term perception of environmental cues translate into the dauer decisions which take place over the course of hours? To address these questions, we studied the dauer exit decision using genetic and molecular neurobiological approaches. Using an ethologically relevant, pheromone-based assay, we demonstrate an essential role for neuropeptide signaling as a whole in dauer exit and validate INS-6 as important for dauer exit.

To understand how INS-6 production relates to sensory perception, we analyzed the spatiotemporal dynamics of *ins-6* expression in response to different environmental cues and found that *ins-6* expression in a pair of chemosensory neurons, the Amphid Single Cilium J (ASJ) neurons, reflects both food and pheromone levels during dauer exit. We found that high *ins-6* expression in the ASJ neurons reflects commitment to exit the dauer state. Through calcium imaging in dauers, we show that sensory integration of food and pheromone can also be seen at the level of sensory neuron activity: ASJ calcium levels increase in response to food, but this response can be suppressed by adding pheromone. We find that *ins-6* upregulation during dauer exit depends on ASJ neuronal activity, cGMP signaling, a CaM-kinase pathway, and is inhibited most potently by the pheromone component ascr#8. Altogether, our data show how the ASJ neurons integrate food and pheromone levels both in the short-term through calcium transients and in the longer term through transcription of *ins-6*, thereby highlighting how neuropeptides can provide the bridge from ephemeral sensory information to sustained physiological changes.

## Results

### A screen of ASJ-enriched neuropeptide genes validates *ins-6* as critical for dauer exit

Neuronal ablation methods using a laser microbeam or transgenic caspases have shown that the chemosensory ASJ neurons are important for dauer exit (Bargmann and Horvitz, 1991; Chung et al., 2013; Cornils et al., 2011), but the precise molecular mechanism by which the ASJ neurons promote dauer exit remains unclear. Having previously shown that neuropeptides are collectively required for dauer entry (Lee et al., 2017), we reasoned that neuropeptides could also be involved in dauer exit. Using a pheromone based dauer exit assay (**Fig. 1B**), we tested five mutants defective in neuropeptide synthesis or secretion and found that four out of the five loss-of-function mutants (*sbt-1/7BT*, *egl-3/PC2*, *egl-21/CPE*, and *unc-31/CAPS*) had significantly lower dauer exit rates than that of wild-type (**Fig. 1C**), strongly indicating that neuropeptide signaling is required for exit. By contrast, *ric-7* loss-of-function mutants, which are impaired for dense core vesicle secretion (Hao et al., 2012), exited dauer at rates higher than that of wild-type, suggesting that RIC-7 may impair the secretion of different neuropeptides than those affected by the other four mutants.

We hypothesized that the ASJ neurons may utilize neuropeptides to promote dauer exit and tested this by screening for defects in dauer exit rates in mutants lacking ASJ-enriched neuropeptides. We analyzed loss-of-function mutants for eleven insulin-like peptide genes, two neuropeptide-like peptide genes, and two FMRF-amide-like peptide genes that were predicted to be enriched in ASJ based on single-cell RNA-sequencing data (Taylor et al., 2021) (**Fig. 1D, Fig. S1A**). Of the fifteen genes tested, *ins-6* mutants showed the most severe dauer exit defect as they exited dauer at the lowest rates compared to wild type. *ins-32* mutants showed a modest dauer exit defect, while mutants defective for *ins-26*, *nlp-80*, *flp-15*, or *flp-34* showed higher than wild-type dauer exit rates, suggesting that the neuropeptides those genes encode may inhibit dauer exit.

### *ins-6* transcription and INS-6 secretion increase quickly during dauer exit

We focused on characterizing the expression pattern of *ins-6* because it showed the most severe impairment in dauer exit of the neuropeptide genes we tested. To characterize how *ins-6* expression responds to environmental improvement during dauer exit, we analyzed *ins-6* transcription and INS-6 secretion using genomically integrated fluorescence reporters that were simultaneously integrated into the same strain (**Fig. 2A; Table S1**). To measure *ins-6* transcription, we constructed *ins-6p::destabilized-YFP (dYFP)*, a transcriptional reporter based off a previous design (Cornils et al., 2011) that fuses both upstream and downstream regulatory regions of *ins-6* to dYFP, a modified YFP variant with a shorter half-life which provides higher temporal resolution (Li et al., 1998), and measured fluorescence intensity in head neurons. To measure INS-6 secretion, we constructed *ins-6::mCherry*, a translational reporter that uses the *ins-6* promoter to drive expression of a fusion between the INS-6 propeptide and mCherry, a protein whose secretion can be tracked using a coelomocyte uptake assay (Speese et al., 2007) in which fluorescence intensity within the coelomocytes reflects the amount of neuropeptide secreted into the body cavity. We built a strain containing simultaneously integrated copies of both these reporters (see **Table S1**) and concurrently measured YFP signal in head neurons and mCherry signal in coelomocytes (**Fig. 2B**).

**Figure 2.**
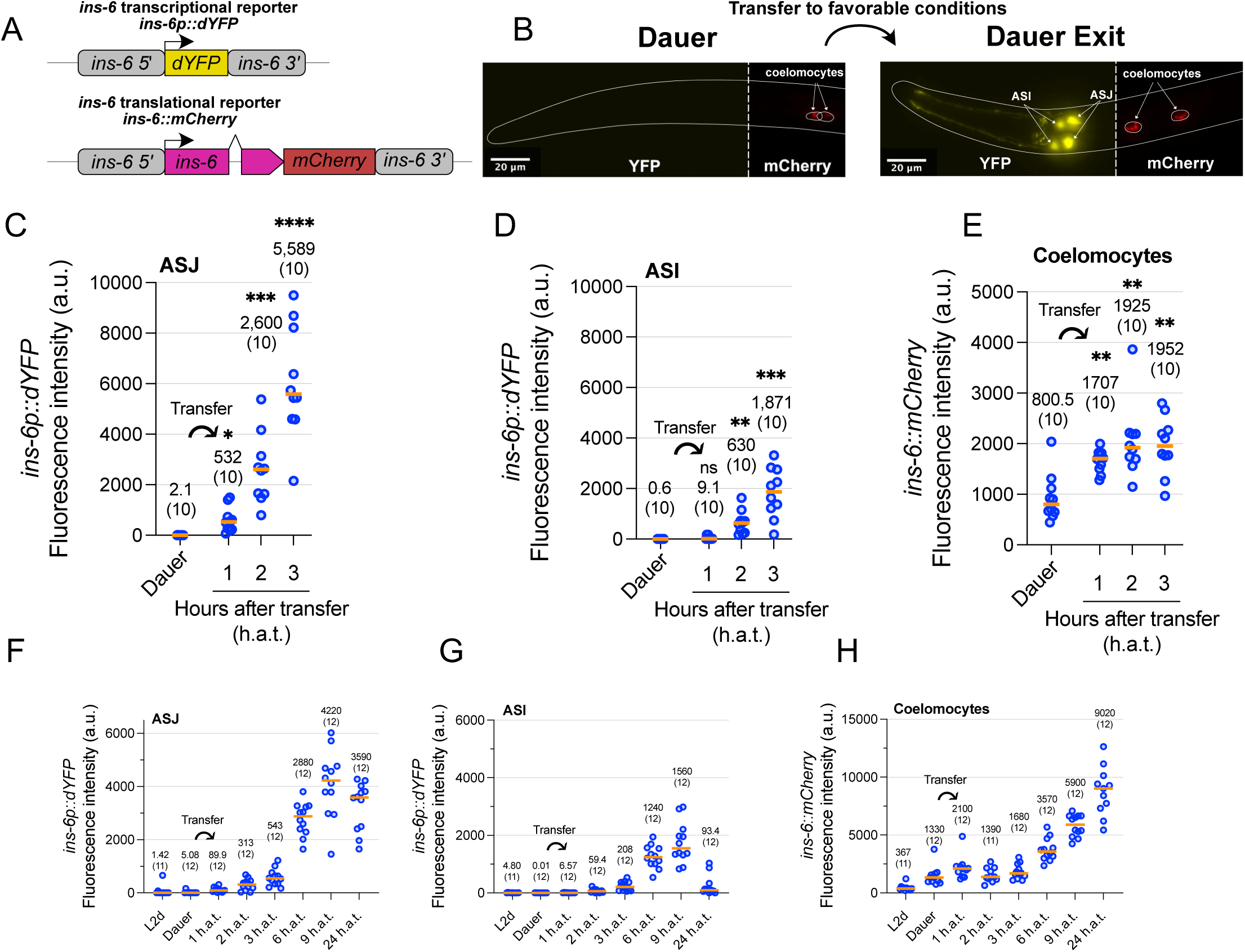
*ins-6* transcription and INS-6 secretion increase quickly during dauer exit. (A) Design of *ins-6* transcriptional and translational reporters. (B) Sample images of merged dYFP and mCherry channels. Left, dauer prior to transfer. Right, dauer six hours after transfer. Anterior is left, ventral is facing the viewer. (C-E) Quantification of *ins-6p::*dYFP transcriptional reporter activity measured in arbitrary units (a.u.) in ASJ (C) or ASI (D) and mCherry signal in the coelomocytes (E) in dauers and following transfer of dauers to favorable conditions (indicated by curved arrow). Plots are from the same set of animals. ns, not significant. *, p<0.05, **, p<0.01, ***, p<0.001, ****, p<0.0001 by Welch ANOVA with Dunnett’s T3 multiple comparison correction when compared to “Dauer”. (F-H) Additional *ins-6* reporter activity experiment performed similarly to (C-E), except with additional time points, and for dYFP measurements, LED power and exposure time were lowered relative to experiments from (C-E) to prevent pixel saturation. h.a.t., hours after transfer to favorable conditions. In all plots, individual dots represent one animal. Medians are depicted by the orange bar. Medians and sample sizes are written.

*ins-6* transcriptional reporter activity in the ASJ neurons of dauers increased in just one hour after transfer of dauers to favorable conditions (**Fig. 2C**) and continued to increase for multiple hours thereafter (**Fig. 2C, F**). *ins-6* transcriptional reporter activity also increased in the ASI chemosensory neurons albeit at a slower rate and to a lower maximum relative to the ASJ neurons (**Fig. 2D, G**). To validate these results, we performed fluorescence *in-situ* hybridization (FISH) for *ins-6* mRNA (**Fig. S2I, J**). Consistent with our fluorescence reporter results, our FISH data indicate that *ins-6* transcripts increased just 1 hour after transfer and continued to increase afterwards. Concordant with results using our *ins-6* transcriptional reporter, we observed that *ins-6::mCherry* translational reporter activity in the coelomocytes increased within the first hour following transfer to favorable conditions and continued to increase for multiple hours thereafter (**Fig. 2E, H**). Under reproductive growth conditions, which result in non-dauer adult development, our *ins-6* transcriptional reporter showed no signal in ASJ throughout development (**Fig. S2A**). *ins-6* transcriptional reporter activity in ASI (**Fig. S2B**) and translational reporter activity (**Fig. S2C**) in the coelomocytes peaked in L1 larvae and then decreased throughout development. These results suggest that favorable conditions cause increased *ins-6* reporter activity specifically in the context of animals in the dauer state. In summary, the transfer of dauers to favorable conditions causes an increase in *ins-6* transcription and INS-6 secretion that promotes dauer exit.

The observed increase in INS-6 secretion (as measured by mCherry signal in the coelomocytes) could result from an increase in *ins-6* transcription, INS-6 processing, INS-6 packaging, and/or dense core vesicle release. To parse these different factors, we built a translational reporter driven by a constitutively active ASJ-specific promoter, *trx-1p::ins-6::mCherry*, and again measured coelomocyte uptake of INS-6::mCherry (**Fig. S2D**). The mCherry signal in coelomocytes increased when dauers were transferred to favorable conditions in a manner similar to animals bearing the *ins-6p::ins-6::mCherry* reporter transgene (fold-change of 2.14x and 4.8x at “6 h.a.t.” versus “Dauer” from two independent lines of the *trx-1p::ins-6::mCherry* reporter compared to 2.68x using the *ins-6p::ins-6::mCherry* translational reporter from **Fig. 2H**), suggesting that secretion of INS-6 is also controlled at the level of dense core vesicle release.

### *daf-28* expression patterns suggest a weaker role in dauer exit versus *ins-6*

The same studies that identified INS-6 as a major regulator of dauer exit also characterized DAF-28, another insulin-like peptide that promotes the reproductive growth trajectory during the dauer entry decision (Li et al., 2003), as a weaker regulator of dauer exit (Cornils et al., 2011; Fernandes de Abreu et al., 2014). Consistent with those observations, loss of *daf-28* gene function did not result in a defect in dauer exit rates in an otherwise wild-type background but did enhance the dauer exit rate defect of *ins-6* loss-of-function mutants (**Fig. 1D, S1B**). To understand how the spatiotemporal regulation of *daf-28* compares with that of *ins-6*, we performed similar transgenic reporter experiments to study *daf-28* (**Fig. S2E-H**).

A *daf-28p::dYFP* transcriptional reporter showed virtually no activity in the ASJ neurons (**Fig. S2F**) of dauers after being transferred to favorable conditions but did show slight activity increase in the ASI neurons (**Fig. S2G**). *daf-28::mCherry* translational reporter activity measured in the coelomocytes of dauers did not increase even 6 hours after transfer to favorable conditions (**Fig. S2H**), unlike the *ins-6::mCherry* translational reporter which increased over two-fold within the first 6 hours (compare with **Fig. 2H**). In contrast, relative to growth under dauer-inducing conditions, growth under non-dauer-inducing conditions resulted in strong *daf-28p::dYFP* transcriptional reporter activity in the ASI neurons (**Fig. S2G**), weaker activity in the ASJ neurons (**Fig. S2F**), and strong activity in the coelomocytes (**Fig. S2H**), with activity for both reporters peaking after animals reached adulthood. Collectively, our transgenic reporter lines for *ins-6* and *daf-28* show inverse expression patterns: *ins-6* transcriptional and translational reporter activity increase during dauer exit but less during reproductive growth, and vice versa for the *daf-28* reporters. These findings align with both ours’ and others’ dauer exit assay results in which INS-6 plays a stronger role in regulating dauer exit than does DAF-28 (Cornils et al., 2011; Fernandes de Abreu et al., 2014).

### *ins-6* expression reflects commitment in the dauer exit decision

*ins-6* expression increases quickly in all dauers when they are transferred to strongly favorable conditions that stimulate 100% of dauers to exit, but what happens when conditions are more ambiguous such that only a fraction of dauers exit? We transferred dauers to a lower pheromone concentration that induces approximately half of dauers to exit (hereby referred to as “intermediate-pheromone conditions”) and measured *ins-6p::dYFP* transcriptional reporter activity (**Fig. 3A**). At three hours after transfer, all worms showed a slight increase in *ins-6p::dYFP* signal, while at six hours after transfer and beyond, a clear bimodality of the population emerges: the worms that would eventually go on to exit dauer (as evidenced by a wider pharynx (**Fig. 3B**), which is a key morphological characteristic of dauer exit (Zhang and Sternberg, 2022)), showed high *ins-6* transcriptional dYFP reporter activity, whereas the worms that should remain as dauers showed a return to baseline levels of dYFP signal, resembling the level of signal seen in dauers.

**Figure 3.**
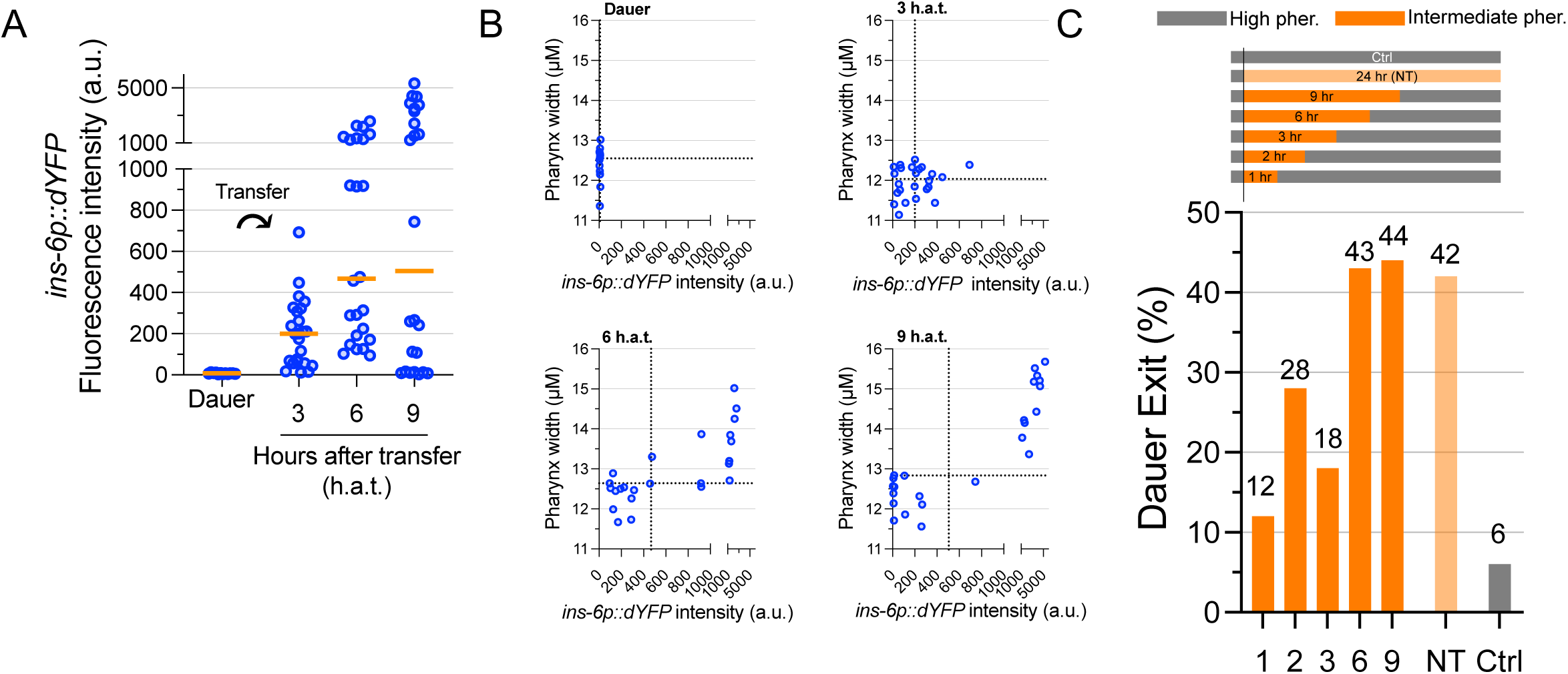
*ins-6* expression correlates with commitment to exit dauer. (A) *ins-6p::dYFP* transcriptional reporter activity in dauers before and after transfer (indicated by curved arrow) onto intermediate-pheromone conditions which stimulate half of dauers to exit. (B) Animals from (A) simultaneously had their pharynx widths measured. Increased pharyngeal width is a key characteristic of ongoing dauer exit. Shown are correlation plots between *ins-6p::dYFP* signal and pharynx width. Dotted lines are drawn at the median for each measurement. (C) Dauers were induced under high-pheromone growth conditions and transferred to intermediate-pheromone conditions for the indicated duration before being transferred back onto high-pheromone conditions to prevent dauer exit in noncommitted animals. Animals were scored for dauer exit after 24 hours. Included are the following controls: a no-transfer control (NT, light orange) in which dauers were not transferred but instead remained on intermediate-pheromone conditions and a control in which dauers remained entirely on high-pheromone conditions (Ctrl, grey). Bars and numbers indicate dauer exit rates for >100 animals per sample.

To determine whether *ins-6* expression could be used to distinguish dauers that are committed versus non-committed, we compared the time course of *ins-6* expression with that of dauer exit commitment (**Fig. 3C**). To examine dauer commitment transferred dauers from high-pheromone, dauer-maintaining conditions to intermediate-pheromone conditions that permitted approximately half the animals to exit dauer, then after different amounts we transferred the animals back onto high pheromone conditions so that animals that had not committed to exiting dauer would be induced to remain as dauers (**Fig. 3C, top**); this protocol was analogous to previous work that established commitment to exiting dauer (Golden and Riddle, 1984). Within three hours of exposure to intermediate-pheromone conditions, approximately half the animals that might exit dauer under such conditions committed to exiting dauer (18% committed, compared to a baseline of 42%), while roughly the total pool of dauers that would eventually exit under intermediate-pheromone conditions committed within six hours (43%) (**Fig. 3C**). These observations indicate that while some dauers irreversibly commit in one hour following transfer to intermediate-pheromone conditions, other dauers can take between three to six hours to commit to the exit decision. This six hour mark matches the time point in our imaging experiments in which the clearest bimodality of both *ins-6p::dYFP* signal as well as pharynx width emerges (**Fig. 3B**), suggesting that such bimodality can reflect decision commitment.

### *ins-6* expression reflects a food:pheromone ratio

Since *ins-6* transcriptional reporter activity responded differently when dauers were transferred to no-pheromone versus intermediate-pheromone conditions (compare **Fig. 2C** to **Fig. 3A**), we asked if *ins-6* expression responds to a food:pheromone ratio – the principal metric by which dauers decide to exit (Golden and Riddle, 1982). Dauers bearing the *ins-6* transcriptional reporter were transferred onto plates containing differing amounts of pheromone and of a food signal (**Fig. 4A)**. We chose crude yeast extract as our food signal because of the following reasons: it promotes dauer exit in our pheromone-based assay (**Fig. S4A**), it is more easily incorporated into agar plates, and because yeast extract is the source material from which a purified food signal was derived in the original dauer exit studies demonstrating the importance of a food:pheromone ratio during dauer exit (Golden and Riddle, 1982).

**Figure 4.**
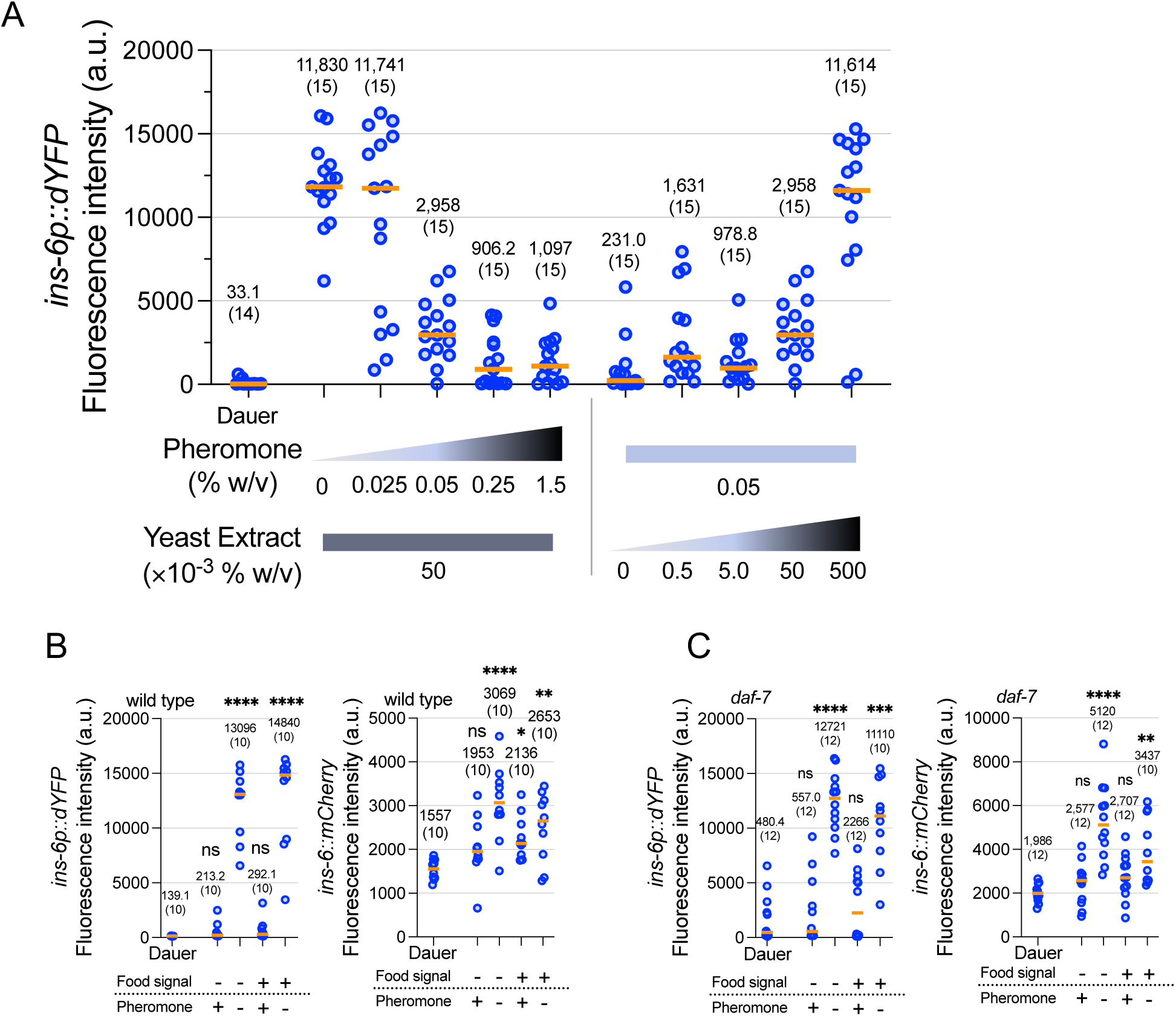
*ins-6* expression reflects food and pheromone integration. (A) *ins-6p::dYFP* transcriptional reporter activity in the ASJ neurons of dauers before transfer (“Dauer”) and three hours after transfer to varying concentrations of food signal (yeast extract) and pheromone. Note that the 0.05% w/v pheromone + 0.05% w/v yeast extract sample is shared between the left and right halves of the graph. (B-C) *ins-6p::dYFP* transcriptional reporter activity in the ASJs (left) and *ins-6::mCherry* translational reporter activity from coelomocytes (right) were measured in wild-type dauers (B) or *daf-7* loss-of-function dauers (which do not exit dauer when kept at high temperatures) (C) before (“Dauer”) and three hours after being transferred to plates with different amounts of food and pheromone (“+”, 0.5% yeast extract or 0.5% pheromone; “-”, 0% yeast extract or 0% pheromone). ns, not significant, *, p<0.05, **, p<0.01, ***, p<0.001, ****, p<0.0001 by Kruskal Wallis Test with Dunn’s multiple comparison correction when compared to “Dauer”. For all graphs: Each dot represents one animal. Medians are depicted by the orange bar. Medians and sample sizes are written.

We found that *ins-6* transcriptional reporter activity correlated positively with food:pheromone ratios (**Fig. 4A**), consistent with *ins-6* being upregulated under favorable conditions during dauer exit. Using live OP50 instead of yeast extract was not more effective in increasing transcriptional reporter activity (**Fig. S4B**). To assess whether food or pheromone was the stronger input, we transferred dauers to plates containing either no pheromone or high amounts of pheromone and either no food signal or high amounts of food signal and analyzed *ins-6* transcription and INS-6 secretion using our transcriptional and translational reporters, respectively (**Fig. 4B**). At high pheromone levels, both transcriptional and translational reporter activity remained low regardless of whether high yeast extract was present, indicating that pheromone can suppress the ability of food signal to increase INS-6 production and secretion.

Having determined that *ins-6* is upregulated under conditions that promote dauer exit, we asked whether this upregulation happens early in the commitment to dauer exit (i.e., happens due to sensory perception of environmental improvement), or whether it might be an effect of the dauer exit process. To explore this possibility, we analyzed our *ins-6* transcriptional and translational reporters in a *daf-7* mutant background. These animals are defective for *daf-7*, which encodes a TGF-β-like growth factor and defines a parallel pathway relative to the insulin-like pathway to which *ins-6* belongs (Hu, 2007; Ren et al., 1996). These mutants constitutively form and remain dauers at 25 °C even under no-pheromone conditions. We repeated the experiment shown in **Fig. 4B** but kept the transferred worms at 25 °C in order to prevent dauer exit. Despite all animals remaining as dauers, these *daf-7* mutants exhibited near-identical *ins-6* regulation with regards to both transcription and secretion when compared to that in a wild-type background (**Fig. 4C**). INS-6 production and secretion is thus upregulated in response to a high food:pheromone ratio that promotes dauer exit, even in animals incapable of exiting dauer, and such upregulation is not simply a downstream effect of dauer exit.

### Calcium imaging in dauers shows integration of food and pheromone in ASJ neurons

To evaluate how the ASJ neurons respond to food and pheromone, we monitored calcium dynamics in dauers expressing the genetically encoded calcium indicator GCaMP6s (Chen et al., 2013) in the ASJs (Hao et al., 2018). We exposed dauers to a food signal (the supernatant from an overnight culture of OP50 grown in LB), pheromone, or a combination of both (**Fig. 5A, S5A**). We found that the ASJ neurons were activated in response to food signal at both stimulus onset and slightly after stimulus offset. The ASJs did not respond to pheromone, but we observed that when the ASJs were exposed to a mixture of food signal and pheromone, the response looked nearly identical to the response to pheromone alone. This finding suggests that ASJ calcium levels integrate food and pheromone. Although the food signal we used differed between calcium and transcriptional reporter imaging experiments, we found that LB (which contains 0.5% w/v yeast extract) increased calcium levels in the ASJs of L4 larvae (**Fig. S5C**), consistent with yeast extract being an activator of ASJ neuronal activity.

**Figure 5.**
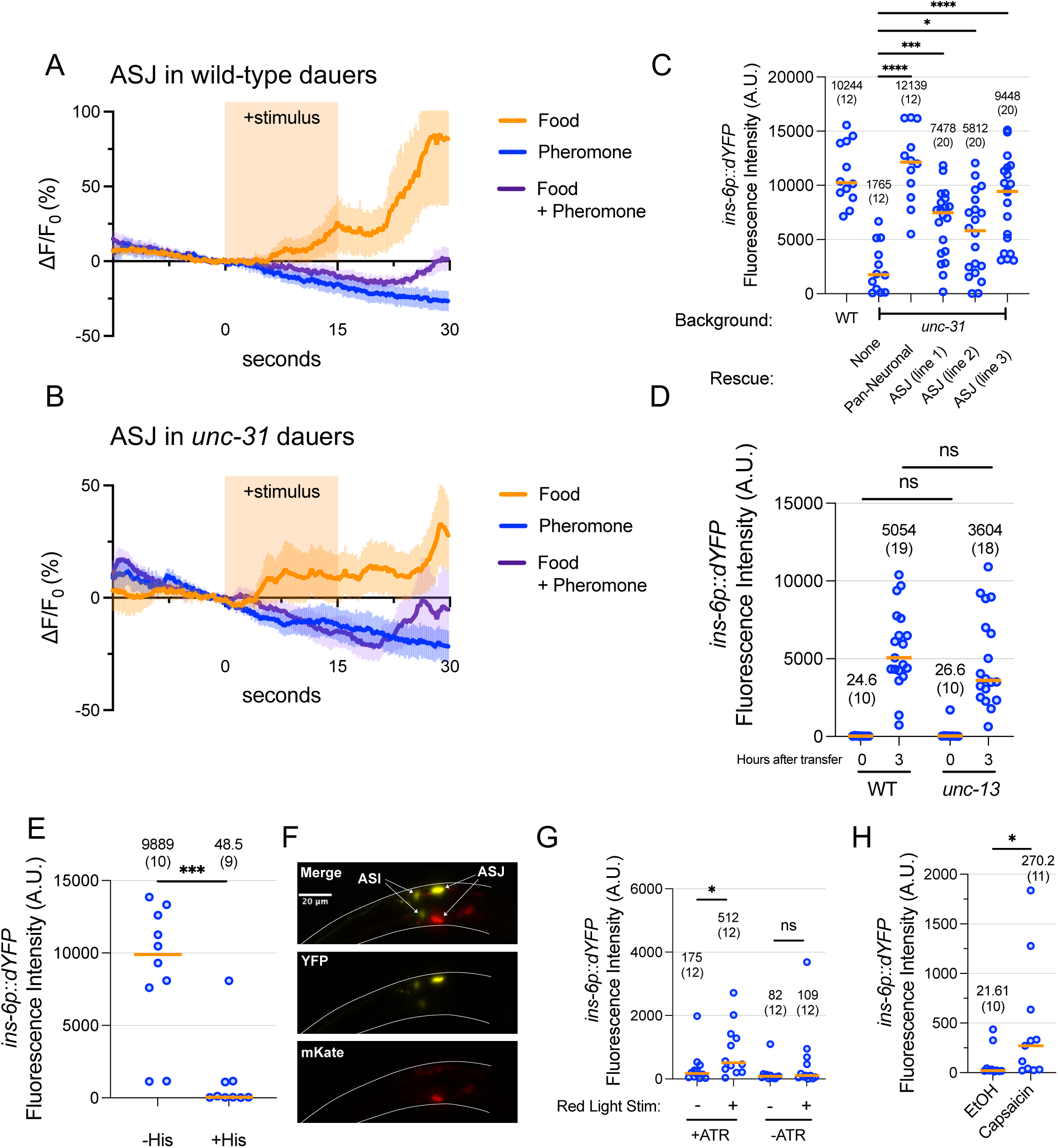
ASJ neuronal activity shows sensory integration and is necessary for *ins-6* upregulation. (A,B) Calcium traces of ASJ in wild-type (A) or *unc-31* loss-of-function mutant (B) dauers in response to pheromone, food (bacterial supernatant), or a mixture of both. Shown are the mean ± SEM from 10-12 ASJ neurons from six animals. Individual traces can be found in Figure S5. (C) *ins-6p::dYFP* transcriptional reporter activity in ASJ was measured in wild-type or *unc-31* loss-of-function mutant dauers (with or without various rescue constructs that express *unc-31* cDNA pan-neuronally or specifically in ASJ) three hours after transfer to favorable conditions. ns, not significant, **, p<0.01, ****, p<0.0001 by one-way ANOVA with Dunnett’s multiple comparisons correction compared to “WT”. (D) *ins-6p::dYFP* transcriptional reporter activity was measured in the ASJs of wild-type or *unc-13* loss-of-function mutant dauers before and three hours after transfer to favorable conditions. ns, not significant, by unpaired t-test. (E) Dauers expressing the histamine-gated chloride channel in the ASJ neurons (*trx-1p::HisCl-T2A-mKate2*) were transferred to favorable plates with or without histamine and imaged three hours later for *ins-6p::dYFP* transcriptional reporter activity in the ASJ neurons. ***, p<0.001 by Mann Whitney U-Test. (F) Image of a mosaic animal in which the *trx-1p::HisCl-T2A-mKate2* extrachromosomal array only expressed in one of the two ASJ nerons. (G, H) Dauers expressing the red-shifted rhodopsin Chrimson (G) or the capsaicin-activated cation channel TRPVI (G) in the ASJs were subjected to the stimulation conditions shown while on high-pheromone conditions and imaged two (F) or three (G) hours later for *ins-6p::dYFP* transcriptional reporter activity in the ASJs. ns, not significant. *, p<0.05 by Mann Whitney U-Test. In all graphs: Each dot represents one animal. Medians are depicted by the orange bar. Medians and sample sizes are written.

### Involvement of other neurons in regulating *ins-6* expression within ASJ

We tested if the *ins-6* regulation to food and pheromone signals in the ASJ neurons depends on signals received from other neurons by measuring *ins-6* transcriptional reporter activity in *unc-31/CAPS* or *unc-13* loss-of-function mutant backgrounds, which are considered defective for neuropeptide secretion and for neurotransmitter release, respectively (Sieburth et al., 2007; Speese et al., 2007). While loss of *unc-31* abrogated *ins-6* upregulation (**Fig. 5C**), loss of *unc-13* did not produce a noticeable effect (**Fig. 5D**), suggesting that neuropeptide signaling, but not fast-acting synaptic transmission, is essential for *ins-6* upregulation. Using an ASJ-specific promoter to express *unc-31* cDNA in the *unc-31* mutant background partially rescued the low levels of *ins-6* transcriptional reporter activity (**Fig. 5C**), implying positive autoregulation: each ASJ neuron might secrete a peptidergic signal to be received by itself or its bilateral partner to increase *ins-6* expression. The lack of a full rescue in two of three independent lines suggests additional neurons may contribute to *ins-6* upregulation by secreting neuropeptides in an *unc-31-*dependent manner. In support of an autoregulatory model, we performed calcium imaging in the ASJ neurons of *unc-31*mutants and found that the responses to food, pheromone, and a mixture of the two remained largely unchanged compared to that of wild type (**Fig. 5B, 5SB**). Specifically, food signal increased calcium levels in the ASJs while pheromone suppressed the positive response to food signal. Collectively, these results imply that when it comes to calcium response, each ASJ neuron possesses the molecular machinery to process both food and pheromone signals and to integrate the two. In addition, the ASJs communicate within a circuit that feeds back to those same ASJs to increase *ins-6* expression in response to food signals.

In accordance with *ins-6* partaking in a feedback loop amongst ASJ neurons, we found that in *daf-2* loss-of-function mutants, which are defective for the insulin/IGF-1 receptor (IGFR) ortholog DAF-2, *ins-6* transcriptional upregulation was impaired (**Fig. S5D**). Transfer of *daf-2* mutant dauers to no-pheromone conditions resulted in a small increase in *ins-6* transcriptional reporter activity compared to that of wild-type dauers (compare **Fig. S5D to Fig. 2C and 4B**). This result is consistent with a model in which *ins-6* regulation depends on an autoregulatory mechanism that involves itself and/or other insulin-like peptides that can signal via DAF-2.

### Activity dependence of ASJ on *ins-6* upregulation

Given the correspondence between ASJ calcium levels and *ins-6* expression when exposed to food and pheromone, we evaluated the relationship between neuronal activity and upregulation of *ins-6* during dauer exit. To test whether *ins-6* upregulation depends on neuronal activity, we chemogenetically silenced the ASJ neurons using cell-specific expression of a histamine-chloride gated channel (Pokala et al., 2014) during dauer exit by transferring dauers to no-pheromone conditions with or without histamine and measuring *ins-6* transcriptional reporter activity. We found that inhibiting ASJ activity severely repressed *ins-6* upregulation (**Fig. 5E**), and that this effect occurs cell autonomously since, in a mosaic animal in which only one ASJ bears the HisCl transgene, only the non-transgenic ASJ showed an increased *ins-6p::dYFP* signal (**Fig. 5F**)

We also tested whether chemogenetic or optogenetic activation of ASJ could induce *ins-6* expression even under high pheromone conditions (**Fig. 5G, H**). Optogenetic stimulation of ASJ via the red-shifted rhodopsin Chrimson (Schild and Glauser, 2015) or capsaicin-induced chemogenetic stimulation of ASJ via the TRPV1 channel (Guo et al., 2015) both slightly increased *ins-6* transcriptional reporter activity, but to a much lesser extent when compared to dauers transferred to favorable conditions (compare **Fig. 5G, H** with **Fig. 2C, 4B**). These loss- and gain-of-function experiments demonstrate that neuronal activity promotes *ins-6* expression during dauer exit.

### Molecular regulation of *ins-6* in ASJ during dauer exit

We sought to characterize the molecular components responsible for *ins-6* upregulation during dauer exit. Since cGMP functions as a messenger in a variety of ASJ-mediated behaviors including dauer development, pathogen avoidance, hydrotaxis, and phototaxis (Birnby et al., 2000; Liu et al., 2010; Park et al., 2020; Wang et al., 2016; Ward et al., 2008), we tested mutants defective for genes in the cGMP pathway. Mutations in *daf-11* (encoding a transmembrane guanylyl cyclase)*, tax-2, or tax-4* (which encode subunits of a cGMP-gated channel) (Coburn et al., 1998; Coburn and Bargmann, 1996) all abrogated *ins-6* transcriptional reporter activity in the ASJ neurons of dauers transferred to favorable conditions (**Fig. 6A**). We also tested the cGMP-dependent kinase EGL-4/PKG (L’Etoile et al., 2002; Stansberry et al., 2001) and found that *egl-4(n479)* loss-of-function mutants had significantly lower *ins-6* transcriptional reporter activity in ASJ (**Fig. 6A**). Addition of the non-hydrolysable cGMP analog, pCPT-cGMP, increased *ins-6* transcriptional reporter activity to high levels even in the presence of pheromone (**Fig. 6B**). Collectively, these results demonstrate that cGMP signaling is both necessary and sufficient for *ins-6* upregulation during dauer exit.

**Figure 6.**
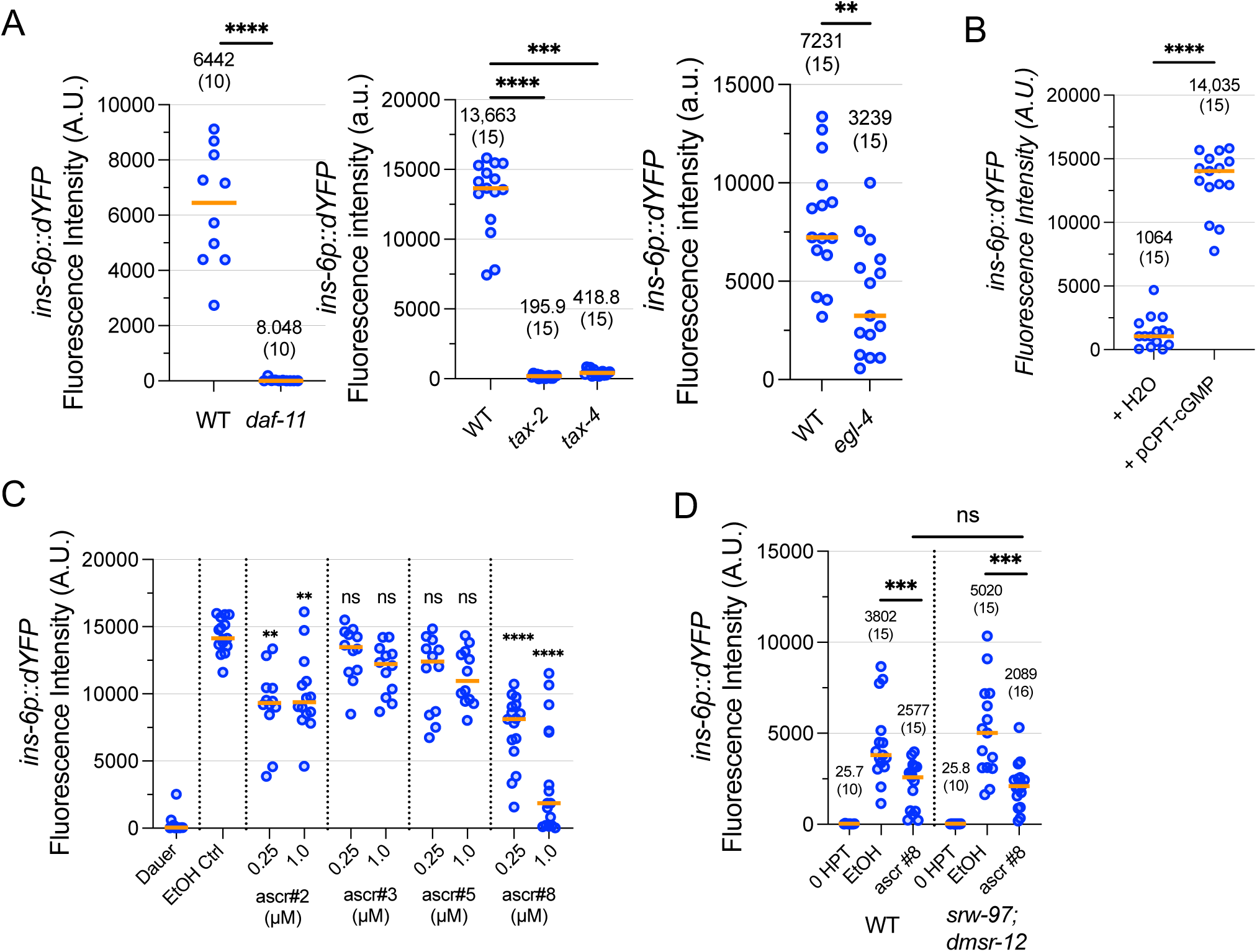
*ins-6* upregulation is regulated by cGMP signaling and ascr#8. (A) *daf-11, tax-2, tax-4,* and *egl-4* loss-of-function mutant dauers were transferred to favorable conditions and measured for *ins-6p::dYFP* transcriptional reporter activity in the ASJ neurons three hours later. ****, p<0.0001 by Mann Whitney U-Test for *daf-11*. ***, p<0.001, ****, p<0.0001 by Kruskal Wallis Test with Dunn’s multiple comparison correction for *tax-2* and *tax-4* compared to “WT”. **, p<0.01 by Mann Whitney U-Test for *egl-4*. (B) Dauers were transferred to high pheromone plates containing 1 mM pCPT-cGMP or H2O control and measured for *ins-6p::dYFP* transcriptional reporter activity in the ASJ neurons three hours later. ****, p<0.0001 by Mann Whitney U-Test (C) Dauers were transferred to plates containing ascarosides at 250 nM or 1 µM and measured for *ins-6p::dYFP* transcriptional reporter activity in the ASJ neurons three hours later. ns, not significant, **, p<0.01, ****, p<0.0001 by Kruskal Wallis Test with Dunn’s multiple comparison correction compared to “EtOH Ctrl”. (C) *srw-97; dmsr-12* loss-of-function mutants were transferred to ascr#8 (1 µM) or EtOH control plates and measured for *ins-6p::dYFP* transcriptional reporter activity in the ASJ neurons three hours later. ns, not significant, ***, p<0.001 by Mann Whitney U-Test. For all graphs: Each dot represents one animal. Medians are depicted by the orange bar. For (A), (B), and (D), medians and sample sizes are written.

To test the role of activity-dependent genes, we measured *ins-6* transcriptional reporter activity in loss-of-function mutants for *cmk-1*, encoding the *C. elegans* homolog of calcium/calmodulin-dependent kinase CaMKI (Eto et al., 1999), and *crh-1*, encoding the *C. elegans* homolog of the CREB transcription factor (Kimura et al., 2002). Both genes have been previously implicated in regulating expression of neuropeptide-encoding genes (Park et al., 2021, 2020; Rojo Romanos et al., 2017). While *cmk-1* mutants displayed a substantial loss in *ins-6* transcriptional reporter activity, *crh-1* mutants showed only a small decrease in reporter activity (**Fig. S6A**). Thus CMK-1 may help transduce calcium influx into upregulation of *ins-6* expression.

Dauer pheromone comprises multiple ascarosides (nematode-specific glycosides of the dideoxy sugar ascarylose) of which ascr#2, ascr#3, ascr#5, and ascr#8 are considered potent inducers of dauer entry (Ludewig and Schroeder, 2018; McGrath and Ruvinsky, 2019). To determine which ascaroside components of the dauer pheromone modulate *ins-6* expression during dauer exit, we transferred dauers to plates supplemented with individual ascarosides at concentrations previously reported to induce dauer entry (Butcher et al., 2008, 2007; Pungaliya et al., 2009) (**Fig. 6C**). We found that ascr#8 most effectively suppressed *ins-6p::dYFP* expression, particularly at 1 µM, while ascr#2 mildly suppressed *ins-6p::dYFP* expression. ascr#3 and ascr#5 did not have a statistically significant effect at either 250 nM or 1 µM. Since ascr#8 most strongly suppressed *ins-6* transcriptional reporter activity in dauers, we tested the GPCRs DMSR-12 and SRW-97, previously shown in *C. elegans* males to be involved in reception of ascr#8 in the context of attraction to hermaphrodites (Reilly et al., 2023), for their roles in regulating *ins-6* expression (**Fig. 6D**). A double mutant defective for both *dmsr-12* and *srw-97* showed a wild-type *ins-6* transcriptional reporter activity response to ascr#8, suggesting additional receptors may be responsible for ascr#8 response in the context of dauer exit.

## Discussion

### *ins-6* highlighted amongst a screen of ASJ-enriched neuropeptides

Having tested fifteen neuropeptide genes enriched in the ASJ chemosensory neurons, we found that *ins-6* loss-of-function mutants had the most severe dauer exit defect in that they exited dauer at the lowest rates compared to wild type (**Fig. 1D**). While *ins-6* has previously been implicated in dauer exit (Cornils et al., 2011; Fernandes de Abreu et al., 2014), this work is the first to test the role of INS-6 function in an otherwise wild-type background for dauer exit. Loss-of-function mutations in *ins-32* also resulted in a strong dauer exit defect, indicating that INS-32 may act alongside INS-6 to promote dauer exit. Loss-of-function mutants for neuropeptide genes such as *ins-26*, *nlp-80*, *flp-15*, and *flp-34* had higher dauer exit rates than those of wild type, suggesting that those neuropeptide genes may be involved in inhibiting dauer exit. Similar to dauer entry, in which we previously demonstrated that some neuropeptides promote while others inhibit dauer entry (Lee et al., 2017), the decision to exit dauer may be partially calculated based on a ratio between exit-promoting versus exit-inhibiting neuropeptides and growth factors.

While the *C. elegans* genome includes over 40 genes encoding insulin-like peptides, *ins-6* stands out as being the only gene whose encoded peptide INS-6 has been shown to biochemically bind and activate the human insulin receptor with high affinity despite bearing key structural differences such as the lack of the conserved C peptide compared to human insulin (Hua et al., 2003; Murphy and Hu, 2018). Such biochemical data provides one possible explanation for the potency of INS-6 in regulating dauer exit. While INS-6 plays a major role in regulating dauer exit, the fact that *ins-6* loss-of-function mutants can still exit dauer under low-pheromone conditions indicates that INS-6 likely acts alongside other insulin-like peptides (such as DAF-28 and possibly INS-32) to induce dauer exit.

### Model for *ins-6* upregulation during dauer exit

We analyzed regulation of *ins-6* using both fluorescence reporter assays (**Fig. 2C-H**) and mRNA FISH (**Fig. S2I, J**), demonstrating that *ins-6* expression and INS-6 secretion increases in just one hour after transfer of dauers onto favorable conditions. In that same time frame, we could not observe any behavioral or morphological changes that would distinguish these dauers as exiting. Our *ins-6* transcriptional reporter showed high activity in the ASJ neurons specifically during dauer exit and virtually no activity when animals were raised under reproductive growth conditions leading to non-dauer adults (**Fig. S2A-C**). These data contrast with previous single-cell RNA sequencing data and mRNA FISH data showing that *ins-6* transcripts are abundant in the ASJ neurons of L4 larvae raised under reproductive growth conditions (Chandra et al., 2019; Taylor et al., 2021). This discrepancy suggests that our *ins-6* reporters contain regulatory regions responsible for dauer-specific transcriptional activity while lacking additional regulatory regions that promote expression in the ASJ neurons during non-dauer-inducing growth.

Our analysis of *ins-6* regulation using fluorescence reporters, gene knockouts, and neuronal perturbations lead us to propose a model of how *ins-6* expression facilitates sensory integration and decision commitment during dauer exit (**Figure 7**). In our model, GPCRs for food signals activate DAF-11 which increases cGMP levels and leads to two parallel outcomes. Firstly, cGMP-gated heterodimeric TAX-2/TAX-4 cation channels open to allow calcium that results in a combination of calcium-dependent transcription mediated through CMK-1/CaMKI as well as exocytosis of neuropeptide-containing dense core vesicles, which has canonically been associated with high calcium levels (Becherer and Rettig, 2006; Ludwig et al., 2002; Ludwig and Leng, 2006; Rizzoli and Betz, 2005; Speese et al., 2007). Secondly, cGMP activates transcriptional activity mediated in part through the cGMP-dependent protein kinase EGL-4/PKG to increase *ins-6* expression. Since neither *cmk-1* nor *egl-4* loss-of-function mutants fully prevented *ins-6* upregulation (**Fig. 6A**), we suspect that both kinases, each defining a calcium-dependent (via CMK-1/CaMKI) or calcium-independent (via EGL-4/PKG) transcriptional pathway, may be necessary for full *ins-6* upregulation during dauer exit. The necessity and sufficiency of cGMP signaling in upregulating *ins-6* during dauer exit may help explain why optogenetic or chemogenetic activation of the ASJ neurons was only sufficient to slightly increase *ins-6* expression under high-pheromone conditions (**Fig. 5G, H**), since calcium alone cannot activate cGMP-dependent pathways such as those involving EGL-4.

**Figure 7.**
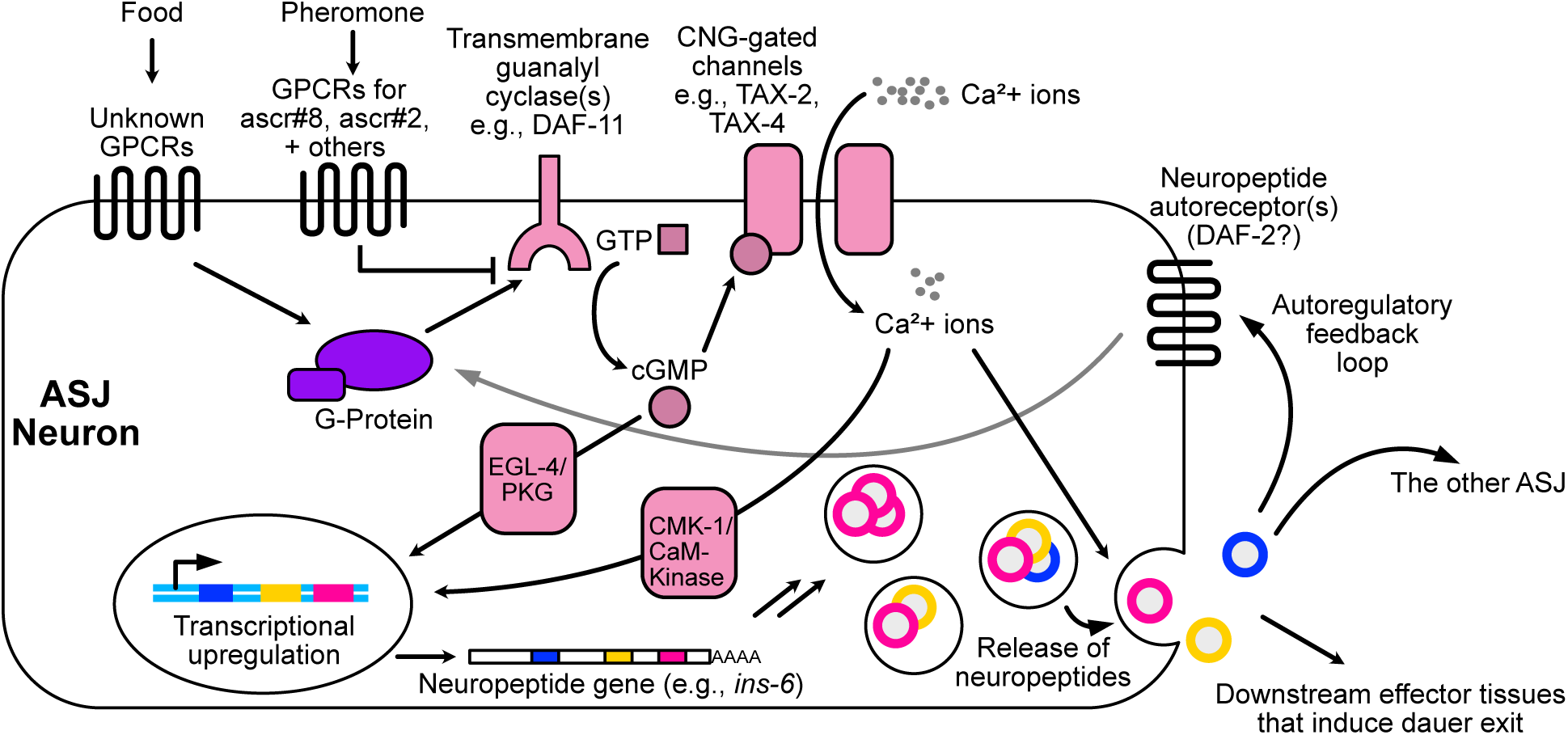
Model for *ins-6* upregulation during dauer exit. In each ASJ neuron, GPCRs specific to food signals trigger a G-protein signaling pathway that activates the transmembrane guanylyl cyclase DAF-11 to increase cGMP production. GPCRs specific to pheromone (including the ascarosides ascr#8 and ascr#2) inhibit this activation. cGMP activates cGMP-dependent transcription pathways such as through the cGMP-dependent kinase EGL-4/PKG while also increasing calcium influx through CNG-gated channels such as TAX-2/TAX-4. The resulting calcium increase can both activate calcium-dependent transcription (such as through CMK-1/CaM-Kinase) and promote secretion of neuropeptides, which may include INS-6, in a process dependent on UNC-31/CAPS. Once secreted, these neuropeptides bind to receptors on the same ASJ and/or the other ASJ neuron and, upon reaching a certain threshold, triggers an autoregulatory positive feedback loop that further increases production and secretion of INS-6 from the ASJs. This positive feedback promotes commitment to exit the dauer state by causing an irreversible buildup of INS-6 production and secretion. INS-6 then signals to downstream tissues that execute the relevant dauer exit gene programs.

How does pheromone inhibit *ins-6* expression? Since pheromone suppressed calcium transients activated by food signal in both wild-type and *unc-31(e169)* loss-of-function mutant dauers (**Fig. 5A, B**), the ASJ neurons likely express GPCRs for pheromone which, upon binding to candidate ligands such as ascr#8 or ascr#2, inhibits calcium influx. Pheromone likely acts upstream of cGMP signaling because addition of pCPT-cGMP increases *ins-6* expression even under high-pheromone conditions (**Fig. 6B**). Therefore, pheromone could prevent calcium influx by blocking food-based activation of DAF-11.

Our observations suggest that *ins-6* expression levels can mark decision commitment. When dauers are transferred to intermediate-pheromone conditions that cause roughly half of dauers to exit within 24 hours, a clear bimodality emerges in *ins-6* transcriptional reporter activity around the same time that dauers commit to exiting (**Fig. 3**). The increase in *ins-6* transcriptional reporter activity in exit-committed dauers likely results from an autoregulatory positive feedback loop that depends on UNC-31 (to help secrete neuropeptides) and DAF-2/IGFR (as the receptor for insulin-like peptides) (**Fig. 5D, Fig. S5D**). We propose that after dauers encounter more favorable conditions, the ASJ neurons produce and secrete small amounts of neuropeptides (which may include INS-6) that bind to receptors (also on the ASJ neurons), thereby triggering an autoregulatory positive feedback loop that promotes further production of INS-6. Our observations that mutating *unc-31* nearly abrogated *ins-6* expression during dauer exit but spared the ASJ calcium response to food and pheromone (**Fig. 5B, D**) suggests that calcium increases are involved upstream of this putative positive feedback loop. Our observation that chemogenetic silencing of the ASJ neurons (which likely inhibits calcium influx) prevents *ins-6* upregulation (**Fig. 5E, F**) further suggests that neuronal calcium may help trigger this positive feedback loop. Such a rampant, irreversible increase in INS-6 production and secretion likely promotes commitment to exit the dauer state by acting on downstream tissues to induce relevant dauer exit-related genetic programs via insulin-like signaling (Murphy and Hu, 2018). Our data that *ins-6* expression increases even in *daf-7* dauers incapable of exiting are consistent with high INS-6 production and secretion as being causes, rather than consequences, of decision commitment.

The nature of this proposed positive feedback loop would explain two features of the dauer exit decision. Firstly, it would explain the all-or-nothing nature of dauer exit in which dauers committed to exiting must proceed fully with decision execution since rampant positive feedback would hinder suppression of that same signaling pathway. Such positive feedback mechanisms have been previously been reported in developmental decisions involving commitment including in yeast, flies, and frogs (Tsuchiya et al., 2014; Xiong and Ferrell, 2003; Yin and Xi, 2018). We previously argued that steroid hormone synthesis may similarly utilize positive feedback to force a binary choice between reproductive growth and dauer during the dauer entry decision (Schaedel et al., 2012). Secondly, the positive feedback loop would help connect the timescale discrepancy between sensory perception (over seconds) and decision execution (over hours). Dauers must be exposed to favorable food:pheromone ratios for a sufficiently long period of time to trigger the level of positive feedback required for decision commitment.

### Ethological significance of *ins-6* expression dynamics in ASJ

How does our model relate to what *C. elegans* may encounter in its natural habitat during dauer exit? In its natural life cycle, *C. elegans* cycles through periods of favorable conditions and unfavorable conditions (boom and bust), and dauer exit plays a critical role in supporting the transition from bust to boom. Expression levels of *ins-6* can convey two important pieces of information: presence of food and absence of pheromone (including ascr#8, which is secreted most by sexually mature hermaphrodites (Kaplan et al., 2011)). We envision that, in the wild, *C. elegans* larvae enter dauer as they grow up in an environment crowded by pheromone-secreting worms and a depleted food source. To exit dauer, *C. elegans* would need to travel away from its crowded environment and find a new food source. At the chemosensation level, this would equate to perceiving that both food is present and that pheromone levels are low. The presence of food alone is insufficient because high pheromone levels indicate that, even if food were present, it would be depleted too quickly given the abundance of other worms. Our results are consistent with this scenario in that high pheromone levels suppress food response at both the calcium and *ins-6* expression level (**Fig. 4B, 5A**).

Our hypothesis for *ins-6* upregulation is consistent with a parallel pathway model that includes a calcium (activity dependent) pathway in addition to a calcium-independent pathway (perhaps mediated through the cGMP-dependent kinase EGL-4/PKG). Such a model has previously been reported for the cases of *daf-7* upregulation (also in the ASJ neurons) in response to *Pseudomonas aeruginosa* and in the case of *flp-19* regulation in the BAG neurons (Park et al., 2020; Rojo Romanos et al., 2017). As postulated previously (Park et al., 2020), having a second, activity-independent pathway would prevent ASJ from inappropriately expressing *ins-6* during exposure to dauer-irrelevant stimuli that activate ASJ including blue light, electric fields, and sodium chloride (Gabel et al., 2007; Ward et al., 2008; Zaslaver et al., 2015), thereby conferring specificity with regards to what environmental cues upregulate *ins-6* in ASJ.

### Sensory integration in a single *C. elegans* sensory neuron

Our genetic and calcium imaging experiments indicate that *ins-6* upregulation during dauer exit occurs cell autonomously within ASJ. *unc-13* loss-of-function mutants defective for synaptic transmission had close to wild-type *ins-6* upregulation in ASJ (**Fig. 5D**), while *unc-31* loss-of-function mutants defective for neuropeptide transmission had low *ins-6* expression levels that could be partially rescued by ASJ-specific expression of *unc-31* cDNA (**Fig. 5C**) but retained wild-type calcium responses to food and pheromone (**Fig. 5B**). Our data suggest that each ASJ neuron alone can integrate food and pheromone at both the level of neuronal calcium and gene expression, which highlights a stark difference between how *C. elegans* integrates sensory inputs compared to insects and vertebrates. The compact nervous system of *C. elegans* requires that it efficiently use its few neurons to discriminate against the vast array of sensory signals. Accordingly, *C. elegans* employs multiple polymodal sensory neurons that express multiple GPCRs to detect different stimuli (Troemel et al., 1995), such as ASH which can detect a variety of noxious stimuli, AWA which responds to numerous chemoattractants (Bargmann, 2006; Ferkey et al., 2021), and AWC which responds to both odors and temperature (Biron et al., 2008; Kuhara et al., 2008). While vertebrates and insects do utilize polymodal neurons, most notably in nociception (Boivin et al., 2023; Emery et al., 2016; Emery and Wood, 2019), sensory integration in such organisms is typically performed via a multilayer model in which modality-specific neurons converge onto higher order brain regions (Ghosh et al., 2017; Yu et al., 2022), a key example being the “one neuron – one receptor” principle observed in mammals and flies wherein each olfactory neuron classes usually only express one olfactory receptor, and signals from each become integrated in further regions of the olfactory bulb (Serizawa et al., 2004; Task et al., 2022). Our results help us better understand how a single neuron performs complex computations of multiple sensory inputs and underscores how the computational power of nematode nervous systems (all about 300 neurons) is vastly underestimated by cell number alone.

## Materials and Methods

### *C. elegans* strains and maintenance

*C. elegans* strains were derived from the wild-type strain N2 (Bristol) and were cultured according to standard laboratory conditions on Nematode Growth Medium (NGM) agar seeded with *E. coli* OP50 as the food source. A list of strains used in this study, including their genotypes and origins, can be found in Table S1. Curated genetic and genomic information was provided by WormBase (Davis et al., 2022).

### Dauer entry induction

To induce for dauer exit and fluorescence imaging assays, 10-20 young adults were placed on 35 mm diameter Petri dishes containing 2 mL of NGM agar (without peptone) supplemented with crude pheromone extract (prepared as previously described; Schroeder and Flatt 2014) at a concentration of 0.5-2.5% w/v, enough to induce >85% animals to enter dauer. Plates were seeded with 10 µL of 8% w/v *E. coli* OP50, which was prepared by pelleting an overnight culture via centrifugation and resuspending in S-Basal three times (hereby referred to as “S-Basal-washed OP50”). Adults were picked onto the plate and allowed to lay eggs at room temperature (RT; 22-23°C) for 5-9 hours before being removed, during which time they typically laid 300-500 eggs. The plates were then further seeded with an additional 50 µL of heat-killed 8% w/v *E. coli* OP50, which was prepared by pelleting an overnight culture via centrifugation, resuspending in S-Basal three times, and placed in a heat block set at 95 °C for 5 minutes (hereby referred to as “heat-killed OP50”). Afterwards, the plates were wrapped with Parafilm (Amcor) and incubated at 25.5-26.1°C for 60-72 hours.

### Dauer exit assay

Dauers were formed according to “Dauer entry induction” (above) and then selected for by an SDS wash (2% in M9, 30 minutes, 25°C) to kill non-dauers before being washed 3x in M9 solution (pelleted by centrifuging for 1 minute at RT, 1000 x *g*). Surviving dauers were then plated onto dauer exit plates seeded with 5 µL 8% w/v heat-killed OP50. Dauer exit plates were prepared similarly to dauer entry plates but contained a lower concentration of crude pheromone extract (typically 0.05% w/v) that had been determined to induce ∼40-80% of wild-type dauers to exit within 24 hours. After 24 hours, dauer exit was scored according to morphological and behavioral criteria, including pharyngeal pumping, body thickening, and paler color (Zhang and Sternberg, 2022). The few experiments in which the wild-type dauer exit rate was not within the 40-80% range were discarded in order to maintain consistency of dynamic range for scoring.

### Molecular cloning and transgenesis

Molecular cloning was performed using NEB HiFi Assembly (New England BioLabs (NEB)) using fragments obtained from PCR with Q5 or Phusion (NEB) or Platinum SuperFi DNA Polymerase (Invitrogen). HiFi Assembly products were transformed into chemically competent DH5α cells via heat shock. Most plasmids used in this study were constructed in a backbone consisting of a derivative of the pSM vector (Shen and Bargmann, 2003), itself a derivative of the Fire vector pPD49.2 (Addgene plasmid # 1686, deposited by Andrew Fire). Plasmids were microinjected into the gonads of young adult animals according to standard protocols (Mello et al., 1991; Rieckher and Tavernarakis, 2017), with most plasmids first linearized via PCR using oMZ15F and oMZ17R to promote cell-specific expression (Etchberger and Hobert, 2008). Contents of injection mixtures can be found in **Table S1**. Select transgenes were integrated according to standard X-Ray irradiation protocols and outcrossed at least three times prior to use (Evans, 2006).

To construct the *ins-6* transcriptional reporter, we took pQZ10 (which includes mCherry flanked by the 1.7 kb upstream and 2 kb downstream of the *ins-6* open reading frame) (Cornils et al., 2011) and replaced the mCherry coding region with that of destabilized YFP, which is a YFP variant that includes a destabilized signal from mouse ornithine decarboxyl at the C terminus that promotes faster degradation (Li et al., 1998). To construct the *ins-6* translational reporter which includes the INS-6 propeptide fused to mCherry at the C terminus, we took pQZ10 and cloned in the full *ins-6* open reading frame (including the sole intron) immediately after the 1.7 kb *ins-6* 5’ upstream fragment. The *daf-28* transcriptional and translational reporters were made in an identical manner using the 3 kb upstream and 2.1 kb downstream of the *daf-28* open reading frame.

### CRISPR Cas9 genome editing

To generate the loss-of-function mutants for *ins-24, ins-32, nlp-21, nlp-80, dmsr-12, srg-36,* and *srx-43*, we used the CRISPR STOP-IN method (Wang et al., 2018). Briefly, C. elegans strain N2 was gene-edited by the insertion of a universal 43-nucleotide-long knock-in cassette (STOP-IN) using CRISPR/Cas9 into an early exon of the target gene to disrupt translation.

### Imaging transgenic *ins-6* reporter strains throughout development

Transgenic animals bearing an integrated fluorescence reporter were semi-synchronized for growth by allowing young adult hermaphrodites to lay eggs for ∼3-5 hours on a 35 mm diameter Petri dish containing 2 mL of NGM agar (without peptone) seeded with 10 µL 8% w/v S-Basal-washed OP50. For experiments involving dauer-inducing conditions, plates contained a high concentration (0.5-2.5% w/v) of crude pheromone and were seeded with 50 µL of 8% w/v heat-killed OP50. For experiments involving non-dauer-inducing conditions, pheromone was omitted, and plates were seeded with 50 µL of 8% w/v S-Basal-washed OP50. Plates were then incubated at 25.5-26.1 °C. For dauer-inducing growth (**Fig. 2C-H**), distinct developmental stages were imaged at the following time points based on the number of hours spent in the incubator: L2d (40-45 hours), dauer (62-66 hours). For non-dauer-inducing growth (**Fig. S2A-C, E-F**): L1 (22 hours), L2 (32 hours), L4 (46 hours), young adult (56 hours), adult (65 hours). To image dauers during dauer exit, dauers were SDS-selected in a manner identical to that used in the “Dauer exit assay” (above) before being transferred to a new NGM agar (without peptone) plate that lacked crude pheromone and was seeded with 5-10 µL of 8% w/v heat-killed OP50. Dauers were then imaged at the indicated time points based on the number of hours after transfer to the new plate.

### Microscopy and image analysis

Animals were immobilized on a pad composed of 4% ultrapure agarose in H2O in 1-2 µL of 5 mM levamisole in H2O or in a suspension of 0.1 µm polystyrene beads (Polysciences) in H2O. Imaging was performed on a Zeiss AxioImager2 equipped with a Colibri 7 for LED fluorescence illumination and an Axiocam 506 Mono camera (Carl Zeiss Inc.). Pharyngeal bulb width measurements were performed using FIJI (ImageJ) software (Schindelin et al., 2012) by using the length tool to measure the widest section of the posterior pharyngeal bulb. Images were processed using FIJI. Quantification of YFP in *ins-6p::dYFP* transgenic animals was performed by drawing an ROI around the neuron of interest (for each animal we measured two ASJ and/or two ASI neurons), measuring the YFP intensity, adjusting for background, and averaging between the two neurons. The ROI was drawn to be approximately the same size and position as that of the neuronal nucleus from the neuron of interest, as estimated by using the DIC channel. Quantification of secreted INS-6::mCherry or DAF-28::mCherry in coelomocytes was performed as described previously (Speese et al., 2007); in short, the four brightest puncta from a maximum intensity projection of the anterior pair of coelomocytes were measured, adjusted for background, and averaged.

### mRNA fluorescence *in-situ* hybridization (FISH)

FISH was performed similarly to a previously used protocol (Chandra et al., 2019). *ins-6* mRNA probes (20 total, see **Table S2** for probe sequences) were designed using the Stellaris RNA-FISH Probe Designer 2.0 (LGC Biosearch Technologies) with non-specificity masking level 5 with the target sequence between the entire *ins-6* open reading frame along with the 5’ and 3’ untranslated regions. Probes were labeled with CAL-Fluor 610 at the 3’ end. At the appropriate developmental stage, animals were washed 3x with M9 and fixed with 4% paraformaldehyde (PFA) in phosphate-buffered saline (PBS) for 45 minutes. Following three additional washes in PBS containing 0.01% Triton X-100, animals were fixed in 100% methanol overnight at 4 °C. The following day, animals were rehydrated in saline sodium citrate (SSC) buffer at 4x concentration (prepared by diluting 20x SSC buffer 5x in H2O) containing 0.01% Triton X-100 (4x SSCT) and then incubated with the FISH probes at 1:500 dilution (final concentration 25 nM) in hybridization buffer (10% w/v dextran sulfate, 15% formamide, 2x SSC) overnight. The following day, animals washed 3x in 10% formamide in 4x SSCT for a total of 1.5 hours at 30 °C, with multiple 4x SSCT washes before and after formamide incubation. Animals were then incubated with DAPI (5 ng/mL in PBS). Following two last 4x SSCT washes, animals were stored and mounted in Vectashield (Vector Labs). FISH imaging was performed on an upright Zeiss LSM880 microscope (Carl Zeiss Inc.) equipped with a 594 nm laser using a Zeiss 63x oil objective. Image quantification was performed using FIJI.

### Calcium imaging

To image dauers, we adapted an existing microfluidic chip design (Schiffer et al. 2021) by incorporating a thinner opening for the worm to accommodate dauers. 1 mM Levamisole was used to semi-immobilize the worms. Each stimulus was delivered to the worm for 15 seconds followed by 30 seconds of H2O. Stimuli were delivered in a random order to account for the effects of memory and adaptation. Fluorescence was recorded with a spinning disc confocal microscope (Dragonfly 200, Andor) and a sCMOS camera (Photometrics Kinetix) that captured fluorescence from GCaMP6.0s (Chen et al., 2013) in the ASJ neurons of KP9672 (see **Table S1**) at 10 ms/.5 μm z-slice, 25 z-slices/volume, and 4 volumes/second. To extract calcium traces, we located the center of each neuronal nucleus and took the average pixel intensities of a 5 μm (X) x 5 μm (Y) x 2.5 μm (Z) (rectangular box around those centers. The neuron-independent background signal was removed and ΔF/F_0_ calculated for each stimulus-response, where F_0_ was the average fluorescence value during the 2.5 seconds before delivery of the stimulus.

### Perturbation assays

For optogenetic assays using the red-shifted channelrhodopsin Chrimson (Klapoetke et al., 2014; Schild and Glauser, 2015), we added all-trans*-*retinal (Sigma-Aldrich) to a final concentration of 500 µM to 35 mm diameter Petri dishes containing 2 mL of NGM agar (without peptone) supplemented with a high concentration (0.15% w/v) of crude pheromone extract to prevent dauer exit. Dauers were transferred onto these plates and placed in the Wormlab Tracking system (MBF Biosciences) where we then turned on a red LED light. The LED stimulus regime was 100 ms on, 1000 ms off at 100% intensity for two hours. Similarly, for chemogenetic assays using TRPV1 (Tobin et al., 2002), we added capsaicin (Sigma-Aldrich) to a final concentration of 100 µM. For cGMP assays, we added pCPT-cGMP (Sigma-Aldrich) to a final concentration of 1 mM.

### Statistical Analysis, Plotting, and Figure Design

Plots were designed and statistical tests were performed using Prism 10.0 (GraphPad). Illustrated figures were designed and drawn using Affinity Designer (Serif).

## Acknowledgments

The TRPV1 plasmid was a kind gift from the Bargmann lab (The Rockefeller University). The pQZ::ins-6 plasmid was a kind gift from the Alcedo Lab (Wayne State University). The strain JSR70 was a kind gift from the Srinivasan lab. Some strains (see **Table S1**) were provided by the National BioResource Project (NBRP) (Yamazaki et al., 2010), particularly from the lab of Shohei Mitani (Tokyo Women’s Medical University Institute for Integrated Medical Sciences), as well as the CGC, which is funded by NIH Office of Research Infrastructure Programs (P40 OD010440). The strain KP9672 *nuIs556[ptrx-1::GCaMP6.0s]* was a kind gift from the Kaplan lab. Technical support was provided by members of the Sternberg lab including Barbara Perry, Stephanie Nava, and Wilber Palma. Microscopy assistance was provided by the Beckman Institute Biological Imaging Facility at Caltech. Critical feedback for the manuscript was provided by Sternberg lab members, particularly Hillel Schwartz and Nicholas Markarian.

## Author Contributions

M.G.Z. and P.W.S. conceived of the study. M.G.Z., performed experiments and analyzed data with help from S.H. and N.F.. M.S. and V.V. designed and performed calcium imaging experiments. H.P. created CRISPR mutants. F.S. provided synthetic ascarosides. M.G.Z. wrote the manuscript with editorial assistance from P.W.S.

