## Supplemental figures and tables for "Sensory integration of food availability and population density during the diapause exit decision involves insulin-like signaling in *Caenorhabditis elegans*"

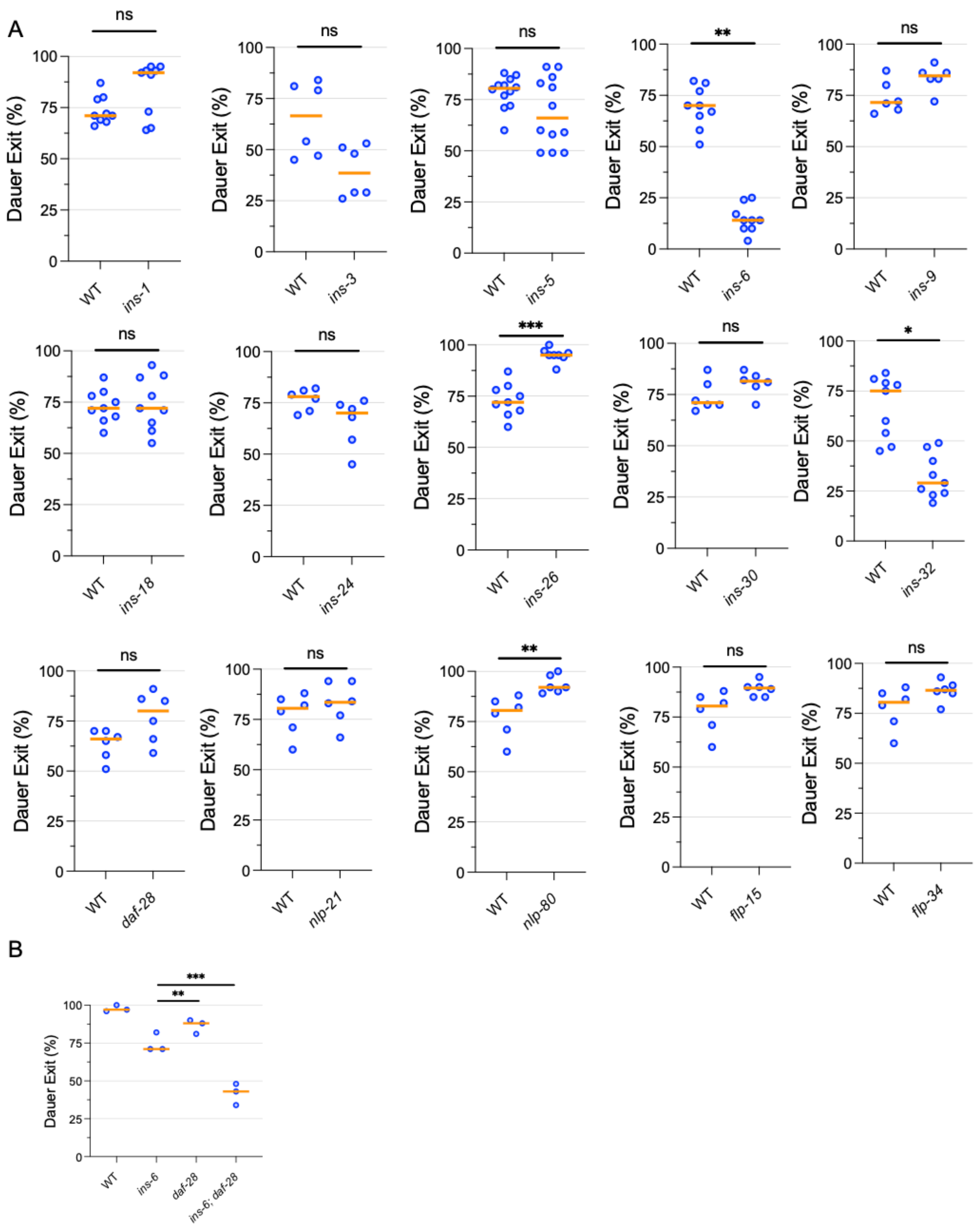


**Figure S1. Supplemental Figure for Figure 1.** (A) Dauer exit rates of mutants defective for single neuropeptide genes. Refer to Table S1 for list of strains. Bars indicate medians. Each dot is the dauer exit % from an assay plate containing 50-100 animals each. ns, not significant, ***, p<0.001, **, p<0.01, compared to "WT" by Welch ANOVA with Dunnett's T3 multiple comparison correction. (B) Dauer exit rates of the double mutant *ins-6; daf-28* compared to single mutants defective for each neuropeptide gene alone. Compared to experiments performed for Fig. 1D, this assay was performed using a lower pheromone concentration that induces more wild-type dauers to exit. Bars indicate medians. Each dot is the dauer exit % from an assay plate containing 50-100 animals each.

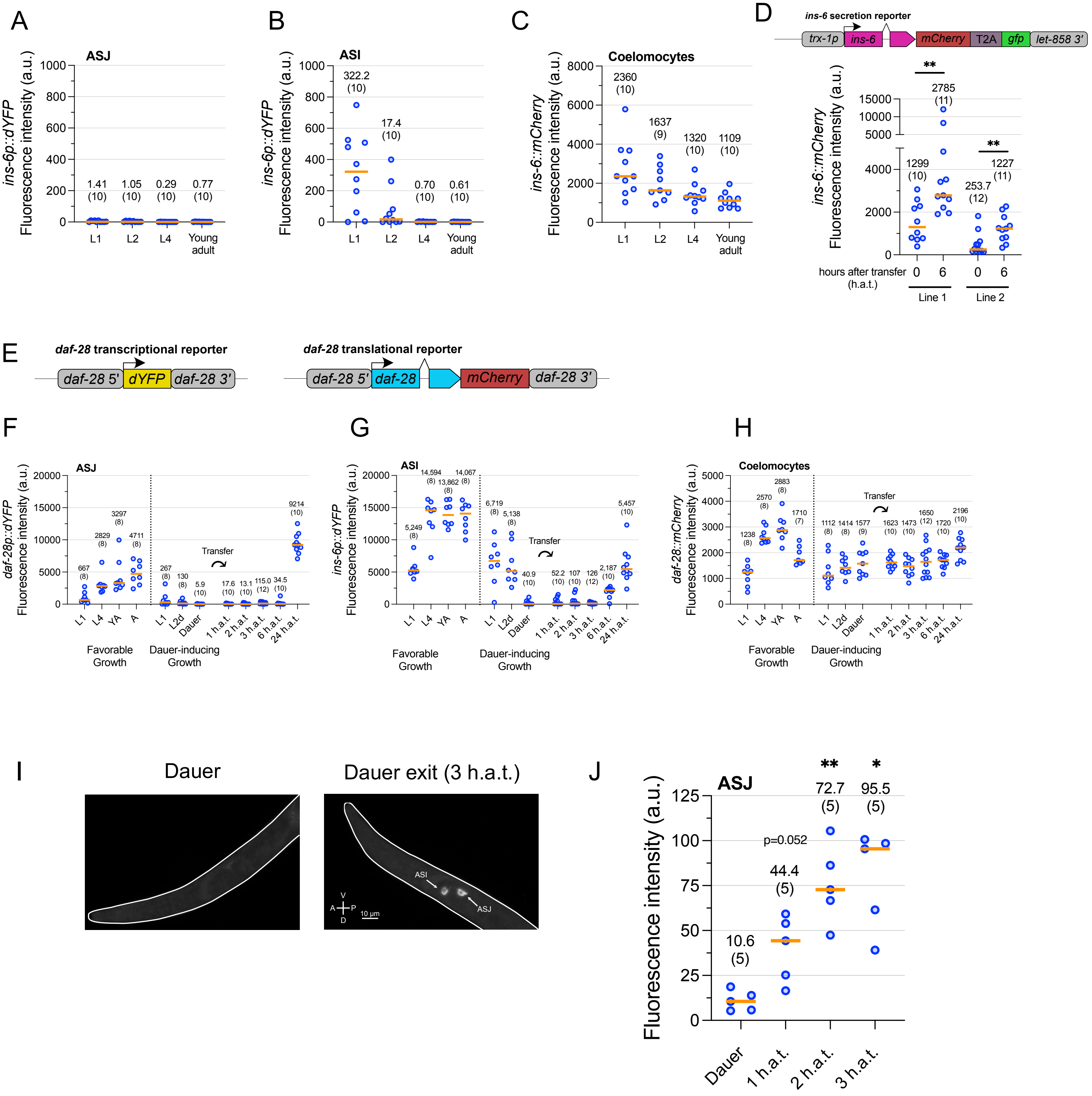


**Figure S2. Supplementary figure for Figure 2.** (A-C) Quantification of *ins-6p::dYFP* transcriptional reporter signal (measured in arbitrary units (a.u.)) in ASJ (A) or ASI (B) and *ins-6::mCherry* translational reporter signal in the coelomocytes (C) during reproductive growth. (D) An *ins-6* secretion reporter was built by driving *ins-6::mCherry* expression using the ASJ-specific promoter *trx-1p* and the broadly expressing *let-858* 3' untranslated region (UTR). Shown are the mCherry fluorescence intensities measured in the coelomocytes for two independent extrachromosomal array lines bearing the *ins-6* secretion reporter in dauers before transfer to favorable conditions (0 h.a.t.) and six hours after transfer to favorable conditions (6 h.a.t.). While the T2A signal was included to simultaneously measure transcriptional activity driven the *trx-1p* promoter, we did not observe any noticeable GFP signal. (E) Design of *daf-28* transcriptional and translational reporters, which were constructed similarly to those for *ins-6*. (F-H) Quantification of *daf-28p::dYFP* transcriptional reporter signal (measured in arbitrary units (a.u.)) in ASJ (F) or ASI (G) and *daf-28::mCherry* translational reporter signal in the coelomocytes (H) during favorable growth (non-dauer-inducing) or dauer-inducing growth, as well as dauers after transfer to favorable conditions (indicated by curved arrow) for the indicated lengths of time. YA, young adult. A, adult. h.a.t., hours after transfer to favorable conditions. See Materials and Methods for precise time points taken during animal development. Dotted lines indicate different populations of animals from which individual animals were sampled for imaging experiments. (I) Representative image of mRNA FISH analysis for *ins-6* mRNA in a dauer (left) and an exiting dauer 3 hours after transfer to favorable conditions). (J) mRNA FISH signal quantification in the ASJ neurons measured in arbitrary units (a.u.). **, $p<0.01$, *, $p<0.05$ by Welch ANOVA with Dunnett's T3 multiple comparison correction when compared to "Dauer". For all quantification graphs, individual dots represent one animal. Medians are depicted by the orange bar. Medians and sample sizes are written.

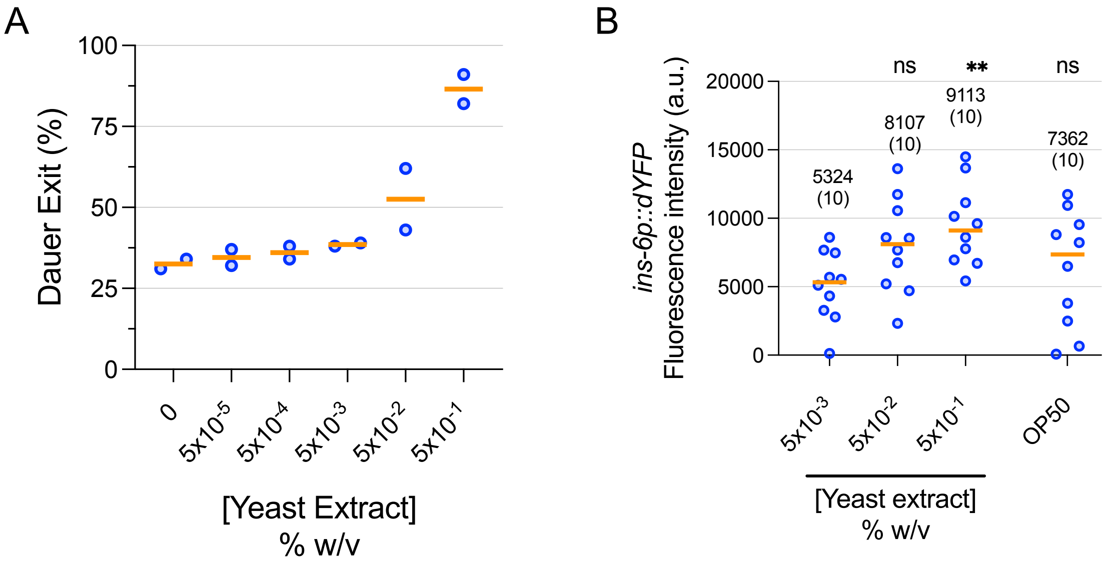


**Figure S4. Supplementary figure for Figure 4.** (A) Dauers were transferred onto plates containing an intermediate-pheromone concentration (0.05% w/v) with varying amounts of yeast extract and scored for exit 24 hours later. Bars indicate medians. Each dot is the dauer exit % from an assay plate containing 50-100 animals each. (B) *ins-6p::dYFP* transcriptional reporter activity was measured in the ASJs of dauers three hours after transfer to plates with an intermediate-pheromone concentration along with various concentrations of yeast extract or a small volume of S-Basal washed live OP50 spotted onto the plate. ns, not significant, **, p<0.01 by Welch ANOVA with Dunnett's T3 multiple comparisons correction when compared to "$5x10^{-3}$% w/v yeast extract". Each dot represents one animal. Medians are depicted by the orange bar. Medians and sample sizes are written.

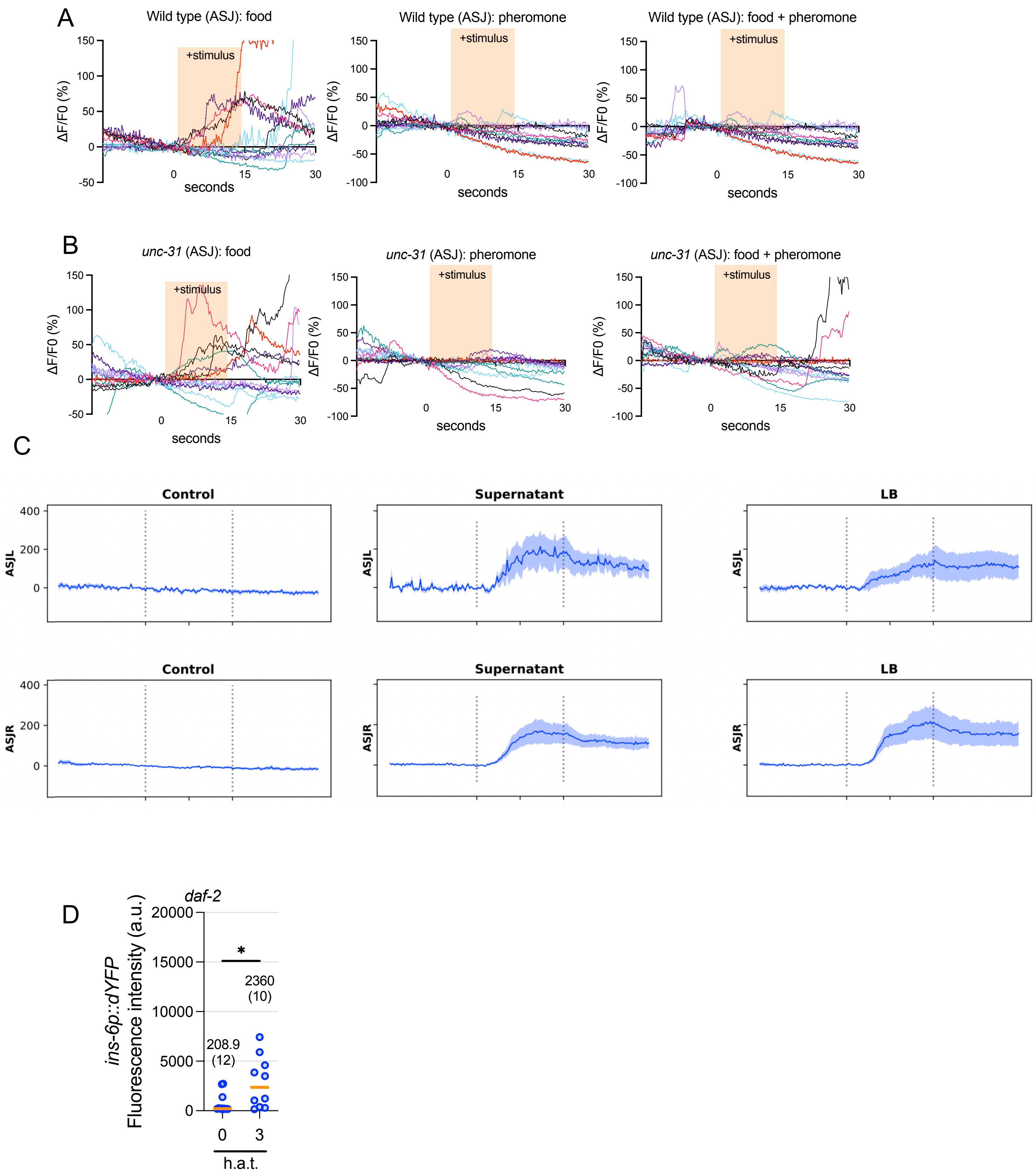

A
Wild type (ASJ): food
Wild type (ASJ): pheromone
Wild type (ASJ): food + pheromone
+stimulus
ΔF/F0 (%)
150
100
50
0
-50
-100
0
15
30
seconds
B
unc-31 (ASJ): food
unc-31 (ASJ): pheromone
unc-31 (ASJ): food + pheromone
+stimulus
ΔF/F0 (%)
seconds
C
Control
Supernatant
LB
ASJL
ASJR
400
200
0
D
daf-2
ins-6p::dYFP
Fluorescence intensity (a.u.)
20000
15000
10000
5000
0
*
2360
(10)
208.9
(12)
0
3
h.a.t.

**Figure S5. Supplemental figure for Figure 5.** (A,B) Individual calcium traces from 10-12 ASJ neurons across six individual animals in wild-type (A) or *unc-31* loss-of-function (B) dauers in response to pheromone, food (bacterial supernatant), or a mixture of both. Aggregate traces are shown in Figure 5A, B. (C) Calcium traces of ASJ neurons (L and R) in wild-type L4 larvae in response to control (buffer), supernatant, and LB. Shown are mean +/- SEM from 7 animals. Each plot includes 15 seconds before stimulus delivery, 15 s of stimulus delivery, and 15 s period after the delivery; the dotted vertical lines indicate the start and end of the stimulus delivery. F0 was defined as average density of the 5 s period before the stimulus delivery. (C) *ins-6p::dYFP* transcriptional reporter activity in the ASJ neurons was measured in *daf-2(e1370)* loss-of-function mutant dauers that were transferred to no-pheromone conditions. Note how the magnitude of *ins-6* transcriptional reporter activity is smaller compared to those of comparable experiments such as in Fig. 2C and 4B. *, $p<0.05$ by Mann Whitney U-Test. Each dot represents one animal. Medians are depicted by the orange bar. Medians and sample sizes are written.

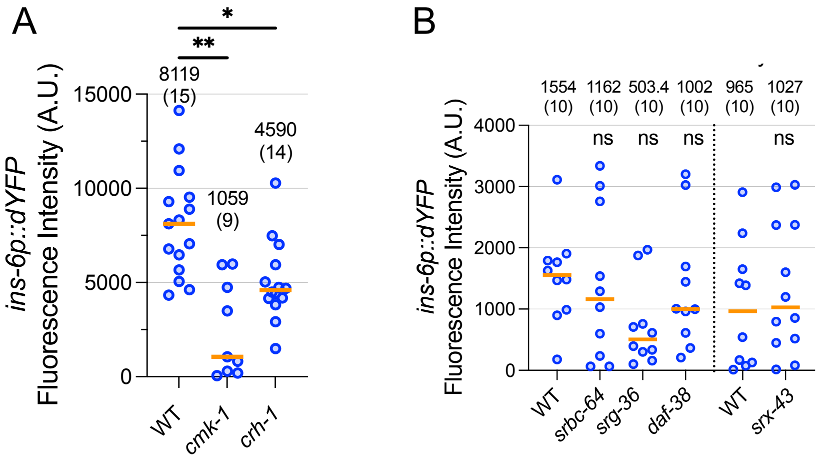


**Figure S6. Supplemental figure for Figure 6.** (A) *cmk-1* and *crh-1* loss-of-function mutant dauers were transferred to favorable conditions and measured for *ins-6p::dYFP* transcriptional reporter activity in the ASJ neurons three hours later. *, $p<0.05$, **, $p<0.01$ by Kruskal Wallis Test with Dunn's multiple comparison correction when compared to "WT". (B) Left of dotted line: *srbc-64*, *srg-36*, and *daf-38* loss-of-function mutant dauers were transferred to threshold pheromone conditions and measured for *ins-6p::dYFP* transcriptional reporter activity in the ASJ neurons three hours later. Right of dotted line: *srx-43(sy1959)* dauers bearing an extrachromosomal array with the *ins-6p::dYFP* reporter transgene were transferred to favorable conditions and measured for YFP signal in the ASJ neurons three hours later. ns, not significant, by Kruskal Wallis Test with Dunn's multiple comparison correction compared to "WT". For all graphs: Each dot represents one animal. Medians are depicted by the orange bar. Medians and sample sizes are written.

**Supplemental Information**

**Table S1: List of strains used in this study**

| Figure | Strain | Genotype | Origin | Notes |
|---|---|---|---|---|
| 1C | N2 | | (Brenner, 1974) | |
| 1C | PS7112 | *sbt-1(ok901)* | (The C. elegans Deletion Mutant Consortium, 2012) | Constructed by outcrossing RB987 to our lab's N2 3x |
| 1C | PS6895 | *egl-3(nr2090)* | (Kass et al., 2001; Liu et al., 1999) | Outcrossed to our lab's N2 4x |
| 1C | KP2018 | *egl-21(n476)* | (Jacob and Kaplan, 2003) | |
| 1C | CB169 | *unc-31(e169)* | (Brenner, 1974) | |
| 1C | KP2048 | *ric-7(nu447)* | (Hao et al., 2012) | |
| 1D | FX01888 | *ins-1(tm1888)* | NBRP (Yamazaki et al., 2010) | |
| 1D | VC1841 | *ins-3(ok2478)* | CGC (The C. elegans Deletion Mutant Consortium, 2012) | |
| 1D | RB2544 | *ins-4(ok3534)* | (The C. elegans Deletion Mutant Consortium, 2012) | |
| 1D | FX2560 | *ins-5(tm2560)* | NBRP (Yamazaki et al., 2010) | |
| 1D | PS9510 | *ins-6(tm2416)* | NBRP (Yamazaki et al., 2010) | Constructed by outcrossing FX2416 to our lab's N2 3x |
| 1D | FX3618 | *ins-9 (tm3618)* | NBRP (Yamazaki et al., 2010) | |
| 1D | VC1218 | *ins-18(ok1672)* | (The C. elegans Deletion Mutant Consortium, 2012) | |
| 1D | PS9417 | *ins-24(sy1761)* | This study | Generated using a CRISPR Stop-In cassette (Wang et al., 2018) |
| 1D | FX1983 | *ins-26(tm1983)* | NBRP (Yamazaki et al., 2010) | |

| | | | | |
|---|---|---|---|---|
| 1D | RB1809 | *ins-30(ok2343)* | (The C. elegans Deletion Mutant Consortium, 2012) | |
| 1D | PS9712 | *ins-32(sy1905)* | This study | Generated using a CRISPR Stop-In cassette |
| 1D | PS9480 | *daf-28(tm2308)* | NBRP (Yamazaki et al., 2010) | Outcrossed ZM7963 to N2 4x and isolated the *daf-28(tm2308)* mutation. |
| 1D | PS9520 | *nlp-21(sy1807)* | This study | Generated using a CRISPR Stop-In cassette |
| 1D | PS8307 | *nlp-80(sy1264)* | This study | Generated using a CRISPR Stop-In cassette |
| 1D | VC2504 | *flp-15(gk1186)* | (The C. elegans Deletion Mutant Consortium, 2012) | |
| 1D | PS7220 | *flp-34 (sy810)* | (Lee et al., 2017) | |
| S1B | N2 | | (Brenner, 1974) | |
| S1B | PS9510 | *ins-6(tm2416)* | NBRP (Yamazaki et al., 2010) | Already described above |
| S1B | PS9480 | *daf-28(tm2308)* | NBRP (Yamazaki et al., 2010) | Outcrossed ZM7963 to N2 4x and isolated the *daf-28(tm2308)* mutation. |
| S1B | PS9518 | *ins-6(tm2416); daf-28(tm2308)* | This study | Cross between PS9480 and FX2416 |
| 2C-H, S2A-C | PS9750 | *syIs875* | This study | *syIs875 [ins-6p::destabilized-YFP; ins-6p::ins-6::mCherry] V* constructed via X-Ray mediated integration of |

| | | | | |
|---|---|---|---|---|
| | | | | *syEx1937[pQZ::ins-6p::dYFP::ins-6UTR (100 ng/uL), pQZ::ins-6::mCherry (25 ng/uL), unc-122p(coel)::gfp (40 ng/uL), 1 kb DNA ladder (35 ng/uL)].* Line 1 of 2 |
| S2D | PS10304 | *syEx1958* | This study | syEx1958[trx-1p::ins-6::mCherry-T2A-gfp::let-858UTR (100 ng/uL), *unc-122p(coel)::gfp* (40 ng/uL), NEB 1 kb ladder (60 ng/uL). Line 1 of 2. |
| S2D | PS10305 | *syEx1959* | This study | syEx1959[trx-1p::ins-6::mCherry-T2A-gfp::let-858UTR (100 ng/uL), *unc-122p(coel)::gfp* (40 ng/uL), NEB 1 kb ladder (60 ng/uL). Line 2 of 2. |
| S2E-H | PS9789 | *syIs876* | This study | Integrated line (via X-Ray irradiation) of PS9725 *syEx1941[daf-28p::dYFP* (100 ng/uL), *daf-28::mCherry* (25 ng/uL), *unc-122p(coel)::gfp* (40 ng/uL), NEB 1 kb ladder (35 ng/uL)] |
| S2I, J | N2 | | (Brenner, 1974) | |
| 3A, B | PS9750 | *syIs875* | This study | Already described above |
| 4A, B | PS9750 | *syIs875* | This study | Already described above |

| | | | | |
|---|---|---|---|---|
| 4C | PS9760 | *daf-7(e1372); syIs875* | This study | |
| S4A-B | PS9750 | *syIs875* | This study | Already described above |
| 5A | KP9672 | *nuIs556[ptrx-1::GCaMP6.0s]* | (Hao et al., 2018) | *nuIs556[ptrx-1::GcaMP6.0s]* |
| 5B | PS10295 | *unc-31(e169); nuIs556* | This study | Cross between CB169 and KP9672 *nuIs556[ptrx-1::GcaMP6.0s]* |
| 5C | PS9903 | *unc-31(e169); syIs875* | This study | Cross between CB169 and PS9750 *syIs875[ins-6p::destabilized-YFP; ins-6p::ins-6::mCherry]* |
| 5C | PS10316 | *unc-31(e169); syIs875; syEx1960* | This study | *syEx1960[*KG#121 *rab-3p::unc-31(cDNA)* 100 ng/uL, *unc-122p(coel)::rfp* 40 ng/uL, NEB 1 kb ladder 10 ng/uL*]* |
| 5C | PS10289 | *unc-31(e169); syIs875; syEx1956* | This study | *syEx1956[trx-1p::unc-31(cDNA)::T2A-mKate2-let858UTR 100 ng/uL, unc-122p(coel)::rfp 40 ng/uL, NEB 1 kb ladder 10 ng/uL] Line 1 of 3* |
| 5C | PS10290 | *unc-31(e169); syIs875; syEx1957* | This study | *syEx1956[trx-1p::unc-31(cDNA)::T2A-mKate2-let858UTR 100 ng/uL, unc-122p(coel)::rfp 40 ng/uL, NEB 1 kb ladder 10 ng/uL] Line 2 of 3* |
| 5C | N/A | *unc-31(e169); syIs875; mzEx116C2* | This study | *mzEx116C2[trx-1p::unc-31(cDNA)::T2A-mKate2-let858UTR 100 ng/uL, unc-122p(coel)::rfp 40* |

| | | | | |
|---|---|---|---|---|
| | | | | *ng/uL, NEB 1 kb ladder 10 ng/uL] Line 3 of 3* |
| 5D | PS9749 | *syIs874* | This study | *syIs874 [ins-6p::destabilized-YFP; ins-6p::ins-6::mCherry] V* constructed via X-Ray mediated integration of *syEx1937[pQZ::ins-6p::dYFP::ins-6UTR (100 ng/uL), pQZ::ins-6::mCherry (25 ng/uL), unc-122p(coel)::gfp (40 ng/uL), 1 kb DNA ladder (35 ng/uL)]*. Line 2 of 2 |
| 5D | PS9851 | *unc-13(e51); syIs874* | This study | Cross between MT7929 *unc-13(e51)* and PS9749 *syIs874 [ins-6p::dYFP; ins-6p::ins-6::mCherry]* |
| 5E, F | PS10284 | *pha-1(e2123ts); syIs875; syEx1954* | This study | *syIs875[ins-6p::destabilized-YFP; ins-6p::ins-6::mCherry] V*<br><br>*syEx1954[trx-1p::HisCl-T2A-mKate2 75 ng/µL, pBX pha-1(+) cDNA 75 ng/µL 50 ng/µL, 1 kb ladder (NEB) 25 ng/µL]* |
| 5G | PS10282 | *pha-1(e2123ts); syIs875; syEx1952* | This study | *syIs875[ins-6p::destabilized-YFP; ins-6p::ins-6::mCherry] V*<br><br>*syEx1952[trx-1p::Chrimson-T2A-mKate2 75 ng/µL, pBX pha-1(+) cDNA 75 ng/µL 50 ng/µL, 1 kb ladder (NEB) 25 ng/µL]* |

| | | | | |
|---|---|---|---|---|
| 5H | PS10330 | *pha-1(e2123ts); syIs875; syEx1964* | This study | *syIs875[ins-6p::destabilized-YFP; ins-6p::ins-6::mCherry] V*<br><br>*syEx1964[trx-1p::TRPV1-T2A-mKate2 75 ng/µL, pBX pha-1(+) cDNA 60 ng/µL 50 ng/µL, 1 kb ladder (NEB) 40 ng/µL]* |
| S5A | KP9672 | *nuIs556[ptrx-1::GCaMP6.0s]* | (Hao et al., 2018) | *nuIs556[ptrx-1::GcaMP6.0s]* |
| S5B | PS10295 | *unc-31(e169); nuIs556* | This study | Cross between CB169 and KP9672 *nuIs556[ptrx-1::GcaMP6.0s]* |
| S5D | PS9759 | *daf-2(e1370); syIs875* | This study | |
| 6A | PS9749 | *syIs874* | This study | Already described above |
| 6A | PS9850 | *syIs874; daf-11(m47)* | This study | Cross between PS9749 *syIs874[ins-6p::destabilized-YFP; ins-6p::ins-6::mCherry] IV* and PS2782 *daf-11(m47) V* |
| 6A | PS9750 | *syIs875* | This study | Already described above |
| 6A | PS10389 | *tax-2(p671); syIs875* | This study | Cross between PS9750 *syIs875 [ins-6p::dYFP; ins-6p::ins-6::mCherry] V* and PR671 *tax-2(p671) I* |
| 6A | PS10390 | *tax-4(p678); syIs875* | This study | Cross between PS9750 *syIs875 [ins-6p::dYFP; ins-6p::ins-6::mCherry] V* and PR678 *tax-4(p678)* |

| | | | | |
|---|---|---|---|---|
| 6A | PS10411 | *egl-4(n479); syIs875* | This study | Cross between PS9750 *syIs875 [ins-6p::dYFP; ins-6p::ins-6::mCherry]* and MT1074 *egl-4(n479)* |
| 6B, C | PS9750 | *syIs875* | This study | Already described above |
| 6D | PS9749 | *syIs874* | This study | Already described above |
| 6D | PS10370 | *srw-97(knu456); syIs874; dmsr-12(sy1559)* | This study | Cross between PS8890 *dmsr-12(sy1959)* (Wang et al., 2018), JSR70 (Reilly et al., 2023), and PS9749. ins-6 reporter in a *srw-97; dmsr*-12 double mutant background. Note that the original *dmsr-12* mutation from JSR70 was exchanged for the new *sy1959* allele created via CRISPR STOP-IN, and the *him-5* mutation was crossed out. |
| S6A | PS9750 | *syIs875* | This study | Already described above |
| S6A | PS10391 | *cmk-1(oy21); syIs875* | This study | Cross between PS9750 *syIs875 [ins-6p::dYFP; ins-6p::ins-6::mCherry]* and PY1589 *cmk-1(oy21)* |
| S6A | PS10396 | *crh-1(tz2); syIs875* | This study | Cross between PS9750 syIs875 [ins-6p::destabilized-YFP; ins-6p::ins-6::mCherry] V and YT17 crh-1(tz2) |
| S6B | PS9941 | *srbc-64(tm1946); syIs875* | This study | Cross between PS9750: *syIs875[ins-6p::dYFP; ins-6::mCherry]* and |

| | | | | |
|---|---|---|---|---|
| | | | | PY6560 *srbc-64(tm1946)* |
| S6B | PS9942 | *syIs875; srg-36(sy1961)* | This study | Cross between PS9750: *syIs875[ins-6p::dYFP; ins-6::mCherry]* and PS9886 *srg-36(sy1961)*, which was generated using a CRISPR Stop-In cassette (Wang et al., 2018) |
| S6B | PS9944 | *daf-38(ok2765); syIs875* | This study | Cross between PS9750 *syIs875[ins-6p::dYFP; ins-6::mCherry]* and PS9943 *daf-38(ok2765),* AKA RB2090 (outcrossed to our lab's N2 3x) |
| S6B | PS9704 | *syEx1937* | This study | *syEx1937[ins-6p::dYFP::ins-6UTR (100 ng/uL), ins-6::mCherry (25 ng/uL), unc-122p(coel)::gfp (40 ng/uL), 1 kb DNA ladder (35 ng/uL)]* |
| S6B | PS10138 | *srx-43(sy1959); syEx1937* | This study | Cross between PS9704 *syEx1937* and PS9884 *srx-43(sy1959)* , which was generated using a CRISPR Stop-In cassette (Wang et al., 2018) |

**Table S2: List of probes used in *ins-6* mRNA FISH**

| Probe # | Sequence |
|---|---|
| 1 | gttgctagatactcgtcaaa |
| 2 | aacggaaaatgagtacgggg |
| 3 | ttggagtgcgcacaaaacga |
| 4 | ttgacggaaacttgcagcga |

| | |
|---|---|
| 5 | ttcagacattgaaggaccga |
| 6 | aagttgcatgcttgctgatt |
| 7 | gttgtgttgaagttcacgga |
| 8 | attggtcggtgagctgattc |
| 9 | tggaacacgtcttgctcgtg |
| 10 | agatgagttttcttccgcag |
| 11 | ccacagacagccatgactaa |
| 12 | cttcttgtgggttgcaaaga |
| 13 | attcagtcgcaatgtccttt |
| 14 | cagaacactgatttccgcag |
| 15 | catggacaacaagcagatct |
| 16 | actgatatcggagtaaaggt |
| 17 | acgagattgtatggtacaga |
| 18 | cacgggtgaaacgagattca |
| 19 | gggatgacagatttgatgag |
| 20 | aggtttattgtacaagccac |
